# The nucleus acts as a ruler tailoring cell responses to spatial constraints

**DOI:** 10.1101/863514

**Authors:** A.J. Lomakin, C.J. Cattin, D. Cuvelier, Z. Alraies, M. Molina, G. Nader, N. Srivastava, J.M. Garcia-Arcos, I.Y. Zhitnyak, A. Bhargava, M.K. Driscoll, E.S. Welf, R. Fiolka, R.J. Petrie, N. Manel, A.M. Lennon-Duménil, D.J. Müller, M. Piel

**Author notes:** These authors contributed equally to the experimental part of this work (A.J. Lomakin, C.J. Cattin). These authors jointly supervised this work (A.J. Lomakin, M. Piel).

## Abstract

The microscopic environment inside a metazoan organism is highly crowded. Whether individual cells can tailor their behavior to the limited space remains unclear. Here, we found that cells measure the degree of spatial confinement using their largest and stiffest organelle, the nucleus. Cell confinement below a resting nucleus size deforms the nucleus, which expands and stretches its envelope. This activates signaling to the actomyosin cortex *via* nuclear envelope stretch-sensitive proteins, upregulating cell contractility. We established that the tailored contractile response constitutes a nuclear ruler-based signaling pathway involved in migratory cell behaviors. Cells rely on the nuclear ruler to modulate the motive force enabling their passage through restrictive pores in complex three-dimensional (3D) environments, a process relevant to cancer cell invasion, immune responses and embryonic development.

**One Sentence Summary:** Nuclear envelope expansion above a threshold triggers a contractile cell response and thus acts as a ruler for the degree of cell deformation.

## Main Text

Much like modern day engineered devices, cells in the human body are able to make measurements. The precision of these measurements plays a critical role in many fundamental physiological and pathophysiological processes. For example, epithelial cells in the intestine monitor local cell densities, and if the density increases above a threshold, the cells exit the tissue, relieving the crowding effect and preventing hyperplasia (*1*). When egressing from the bone marrow or encountering the porous interstitial collagen matrix, hematopoietic cells estimate the pore size of the barrier tissue to choose the site of least mechanical resistance (*2, 3*). Epidermal stem cells sense the amount of the extracellular matrix (ECM) that is available for cell attachment and spreading, and use this as a guidance cue in their cell fate decision-making (*4*). These examples illustrate the sensitivity of complex cell behaviors to environmental spatial and mechanical constraints, known in quantitative sciences as boundary conditions (BCs) (*5*). Although the importance of BCs in cell physiology is increasingly recognized, only a few mechanisms by which cells can measure specific BCs are precisely identified (e.g. the stiffness of the substrate on which cells grow (*6*), or the geometry of their adhesive environment (*7*)). Amongst the known mechanisms, most are related to either strain (deformation) or stress (forces) and are collectively referred to as mechanotransduction pathways (*8, 9*).

Here, we asked whether cells are also equipped with a mechanism to measure absolute dimensions, which could instruct them about distances between neighboring cells or matrix pore size. In our previous study, we discovered that many histologically unrelated cell types change their migratory strategies in response to the specific confinement height (*10*). This almost universally leads to a long-lasting increase in actomyosin contractility and amoeboid cell propulsion in the absence of specific adhesion to the substrate. Together with similar findings in early zebrafish embryos (*11*), these observations illustrate the simplest case in which cells measure one of their dimensions to adapt their behavior to local BCs *in vitro* and *in vivo*. However, the mechanism underlying this phenomenon remained unknown. To tackle this question, we applied a reductionist approach in which the degree of confinement is precisely controlled and paralleled with quantitative microscopy. We confined single nonadherent, initially rounded, interphase cells using an ion beam-sculpted flat silicon microcantilever (Fig. 1A) mounted on an atomic force microscopy (AFM) setup (*12–14*) and simultaneously monitored the actomyosin cytoskeleton dynamics and contractile force generation employing confocal videomicroscopy and AFM-based force spectroscopy.

**Figure 1:**
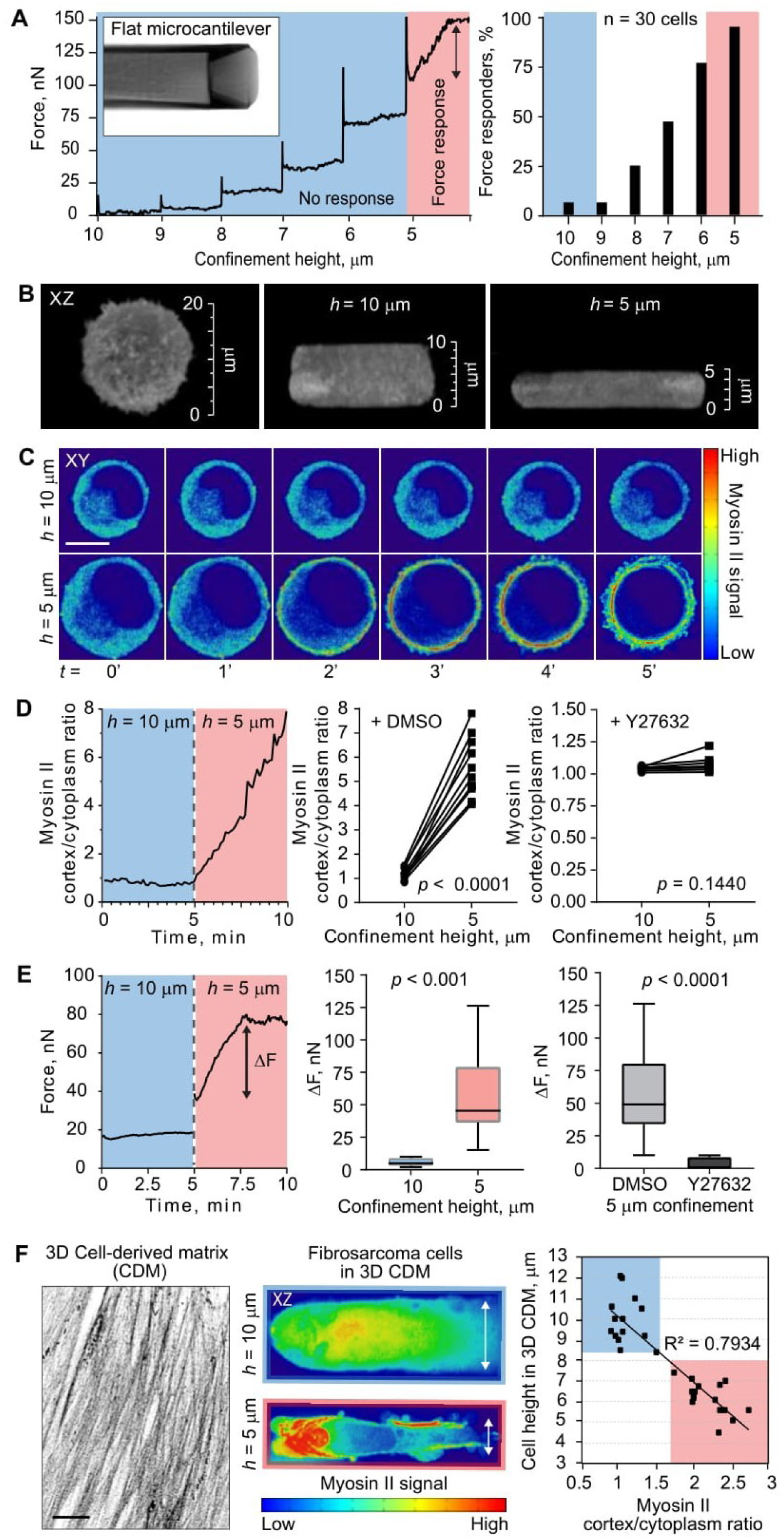
Cells sense their own height and upregulate actomyosin contractility once the height is decreased below a specific threshold. **A:** Left, an XY view of the flat silicon microcantilever and a representative force response curve of a cell whose height was changed in a step-wise fashion from 10 to 5 μm with 1 μm increment and 5 min time interval between each step. Right, percentage of cells displaying the force response (a sustained force increase (> 15 nN)) upon subjecting cells to a specific height. **B:** Cell height changes imposed by the flat silicon microcantilever. 3D images (XZ views) of the same single live HeLa-Kyoto cell expressing Myh9-GFP at 20, 10, and 5 μm confinement height. **C:** Time-lapse image sequence of the same single live cell (XY views) subjected to 10 μm confinement (top) followed by height change to 5 μm (bottom). Each imposed confinement height was stably maintained for 5 minutes and myosin II subcellular distribution over time was monitored simultaneously. Scale bar, 10 μm. **D:** Left, a representative graph illustrating dynamic relocalization of myosin II from the cytoplasm to the cell cortex in the same single live cell confined to 10 μm followed by height change to 5 μm. Middle, change in myosin II cortex/cytoplasm ratio measured in the same single live cell confined to 10 μm and 5 μm (n = 10 different cells; *p* value, paired t test). Right, cancellation of myosin II accumulation at the cortex revealed in the same single live cell (n = 10 different cells; *p* value, paired t test) confined to 10 and 5 μm upon pharmacological suppression of myosin II activity with the Rho kinase/ROCK inhibitor Y27632. **E:** Left, a representative force response curve of the same single live cell confined to 10 μm followed by height change to 5 μm (ΔF, force increase). Middle, statistical analysis of force response output detected at 10 *vs*. 5 μm confinement (mean ± SD; n = 10 cells per each height; *p* value, unpaired t test). Right, statistical analysis of force change in response to 5 μm confinement in cells treated with DMSO (control) and Y27632 (myosin II activity downregulation) (mean ± SD; n = 10 cells per each height; *p* value, unpaired t test). **F:** Left, 3D dermal fibroblast cell-derived matrix (CDM) stained with a collagen I antibody. Middle, representative images of human fibrosarcoma cells HT1080 expressing GFP-myosin light chain 2 in 3D CDM (XZ views, vertical cross-sections) with 10 and 5 μm height. Right, correlation between natural (as opposed to imposed in AFM experiments) cell height in 3D CDM and cortical accumulation of myosin II (n = 30 cells). Scale bar, 20 μm.

Using the cell line HeLa-Kyoto (human cervical carcinoma) expressing MYH9-GFP (Myosin IIA), we performed confinement experiments in which the height of the same single cell (n = 30 different cells) is changed in a step-wise fashion starting from 20 μm (the average diameter of these cells is 20 ± 4 μm in the nonadherent rounded state that we start from in these experiments, n = 100 cells). In a typical experiment, force recording showed that each confinement step is accompanied by an abrupt, large-amplitude increase in output force immediately followed by a flat plateau, which is expected when deforming a passive viscoelastic object (Fig. 1A). However, each analyzed cell at a specific height displayed a steady increase of the force (6 to 5 μm confinement in Fig. 1A) instead of the flat plateau. This steady force increase represents an active response of the cell to spatial confinement and means that the cell is actively pushing the confining cantilever up. Within the cell population, different cells have their own trigger height at which they start generating the outward force on the cantilever (Fig. 1A right graph). Most cells remain insensitive to 10 μm confinement (which corresponds to half of the initial cell height, Fig. 1B), while almost 100% of analyzed cells display an active force response upon reaching 5 μm confinement (Fig. 1A right graph). We thus chose to systematically study the response of cells to 5 *vs*. 10 μm confinement height.

Our analyses showed that cell confinement to 5 but not 10 μm stimulates rapid (2.05 ± 0.33 min, n = 10 cells) recruitment of myosin II from the cytosol to the cortex (Fig. 1C,D and Movie S1), which is then followed by circumferential contraction of the cell and force production (Fig. 1E). Both phenomena require myosin II activity, as follows from the effect of pharmacological inhibition of the upstream regulator of myosin II activity Rho-associated protein kinase ROCK (Fig. 1D,E). The mechanochemical changes observed in response to the threshold confinement manifest themselves in a sustained (up to several hours) and active non-apoptotic plasma membrane blebbing (Movie S2), whose degree is directly proportional to the levels of myosin II at the cell cortex (Fig. S1A-C). Quantifying cell blebbing index as a simple morphological readout of increased actomyosin contractility in several other primary and immortalized cell lines, we were able to confirm the generality of our observations (Fig. S1D). To test whether cells would also adapt their cortical actomyosin contractility to the degree of environmental confinement in a context more closely recapitulating *in vivo* settings, we examined human fibrosarcoma cells HT1080 infiltrating 3D cell-derived matrices (CDM) (Fig. 1F). We found that cortical recruitment of myosin II in these cells linearly scales with self-imposed cell heights, which the cells acquire spontaneously squeezing through differentially sized gaps and trails in the 3D CDM (Fig. 1F), thus validating our AFM-based observations in physiologically relevant conditions. Importantly, switching cell height back to the initial unconfined state in our AFM experiments induced a rapid (3.78 ± 0.94 min, n = 7 cells) re-localization of myosin II to the cytosol (Fig. S1E), indicating that persistent contractility requires a sustained confinement below the threshold height. Collectively, these experiments show that single cells can sense the difference between 10 and 5 μm and trigger a sustained, yet reversible, active contractile response at a specific deformation height.

We next performed experiments to narrow down the range of potential mechanisms involved in this height-dependent actomyosin contractility response. Confinement experiments on adherent, well-spread cells showed qualitatively the same threshold-like response as we established for rounded nonadherent cells (Fig. S2A-C). Moreover, experimental manipulations of extracellular [Ca^2+^] or [Mn^2+^] to modulate engagement of integrins during cell contact with the surface of the confining cantilever (*15*) did not affect the response in nonadherent suspended cells (data not shown). This suggested that the sensing mechanism does not depend on classical integrin-based mechanotransduction pathways. The sustained increase in contractility (Fig. S1C), and the fact that the response was dependent on the confinement height *per se* rather than the speed of confinement (Fig. S3), rules out a signal originating from strain in the actin cortex or the plasma membrane, because these structures dissipate stress in minutes due to fast turnover (*16*). A natural candidate that matches the range of relevant confinement heights at which the response is triggered and that can display long-term stress due to slow turnover of its stiff elastic shell is the cell nucleus (*17*). Indeed, the nucleus, and more specifically its envelope, has been shown in the recent years to trigger diverse cell responses when the nuclear compartment is deformed: entry of the transcription factors YAP/TAZ (*9*), activation of the ATR kinase (*18*), release of calcium (*19*), activation of the phospholipase cPLA2 (*20*) and nuclear envelope rupture accompanied by DNA damage (*21, 22*). Considering that the response to confinement was reversible and required only a few minutes for its manifestation, potential changes at the level of cell transcription/translation are unlikely at play, which we confirmed experimentally by acutely inhibiting the processes of transcription and translation (Fig. S4A). Blocking the ATR kinase activity did not yield a phenotype either (Fig. S4B). Moreover, neither nuclear envelope ruptures nor ruptures of the plasma membrane were observed at 5 μm confinement (Fig. S4C,D), excluding a mechanism based on an extracellular signal influx through transient holes in the plasma membrane, or mixing of cytoplasmic and nuclear contents.

We thus performed a small pharmacological inhibitors screen targeting the pathways compatible with a minute timescale response (e.g., intracellular calcium release – 2APB and BAPTA-AM treatments – and/or signaling lipid production by the calcium-dependent nuclear membrane stretch-sensitive phospholipase cPLA2 – AA (AACOCF3) treatment). We also included inhibition of the myosin light chain kinase MLCK (ML7 treatment) that upregulates myosin II in response to calcium release (*23*), as well as known inhibitors for classical mechanotransduction pathways triggered by plasma membrane tension/cell cortex deformation (see Table S1). The screen (Fig. 2A) showed that extracellular calcium, plasma membrane-associated stretch-sensitive channels and plasma membrane tension are not involved in the response to confinement. However, intracellular calcium, intracellular stretch-sensitive calcium channels associated with the perinuclear ER, the calcium-dependent myosin kinase MLCK and the nuclear envelope tension sensor cPLA2 were required for the contractile response, pointing to a signal emanating from the perinuclear ER and/or the nuclear envelope to activate actomyosin contraction at a specific confinement height. Consistently, imaging of intracellular calcium using the GCaMP6 calcium biosensor revealed a strong increase in cytosolic calcium upon 5 μm confinement (Fig. S5A), which was inhibited by blocking intracellular stretch-sensitive calcium channels InsP3Rs with 2APB (Fig. S5B), but not *via* chelating extracellular calcium with BAPTA (Fig. S5A). Conversely, adding ionomycin (to artificially increase cytosolic calcium concentration) or the signaling lipid arachidonic acid/ARA (the product of active cPLA2 enzyme) to cells confined at 10 μm induced persistent blebbing without further confinement (Fig. S5C). Finally, analysis of the supernatant of a population of confined cells (using a large confinement device, see Methods section) showed an increase in the presence of ARA upon 5 μm confinement, which is lost upon cPLA2 inhibition (Fig. S5D). Importantly, unlike inhibitors that globally perturb basal cell contractility (e.g. Y27632) at both 10 and 5 μm, the drugs that yielded a phenotype in our mini-screen exert their effect only at 5 μm, as follows from our measurements of cortical cell tension and cell pressure at 10 μm (Fig. S5E). This indicates that the targets of the drugs become functionally engaged in a trigger-like fashion when the cell is confined below 10 μm. These results, together with the controls for drugs’ activity (Fig. S5F) suggest that the nuclear membrane compartment, continuous with the perinuclear ER, might be involved in measuring the cell dimensions and triggering the contractile response below a specific confinement height.

**Figure 2:**
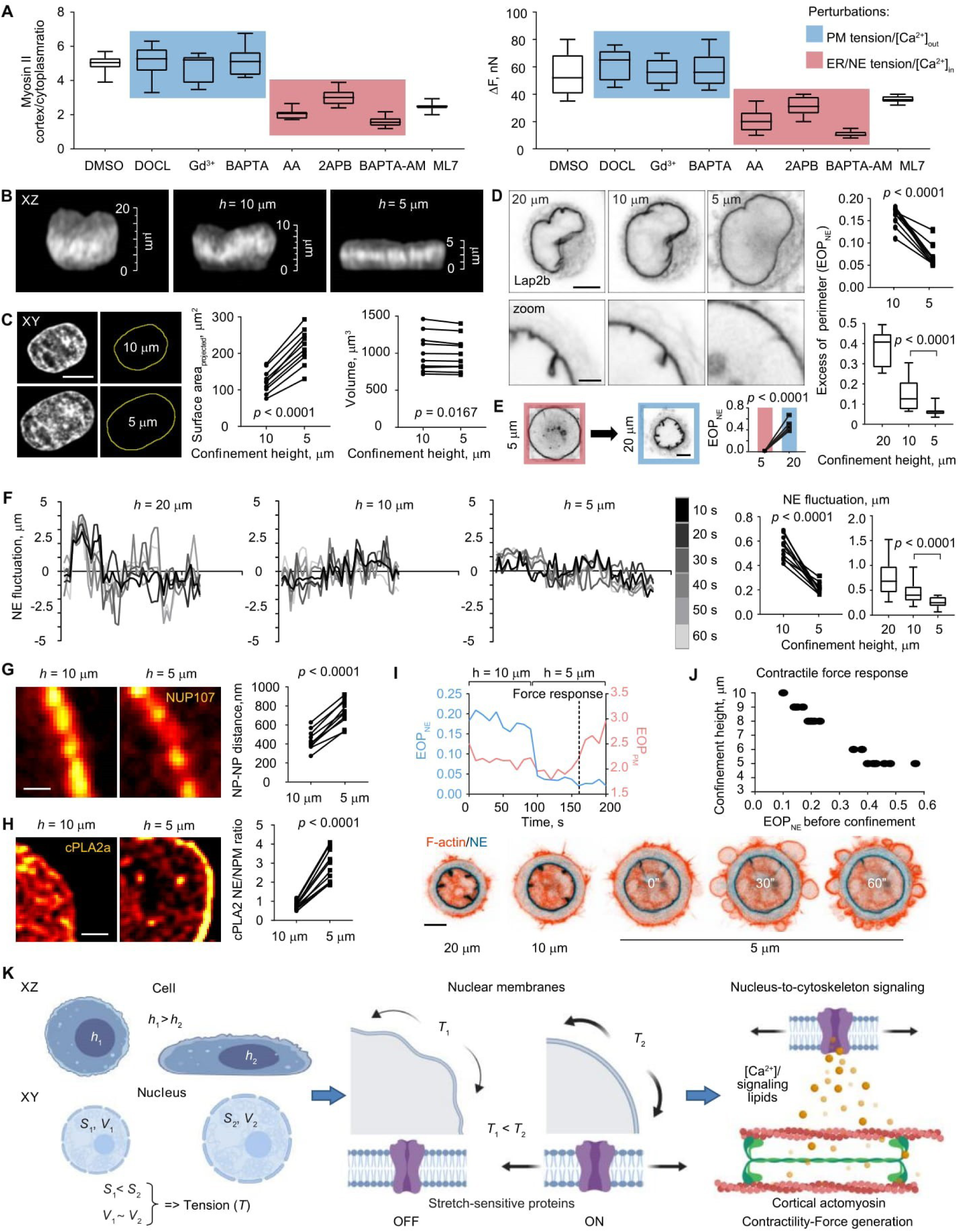
Height-dependent actomyosin contractility output is controlled by the molecular and biophysical mechanisms associated with nuclear and perinuclear ER membrane stretch. **A:** Statistical analysis of myosin II accumulation at the cortex (left) and force generation (ΔF, right) in response to 5 μm confinement in cells treated with DMSO (control), DOCL (deoxycholate, decreasing plasma membrane tension), Gd^3+^ (GdCl_3_, inhibiting stretch-sensitive ion channels on the plasma membrane), BAPTA (an extracellular Ca^2+^ chelator), AA (AACOCF3, blocking the activity of the nuclear envelope tension sensor cPLA2), 2APB (an inhibitor of the stretch sensitive Ca^2+^-channels InsP3R present on the perinuclear ER and nuclear membranes), BAPTA-AM (an intracellular Ca^2+^ chelator), and ML-7 (an inhibitor of the Ca^2+^-calmodulin-dependent myosin light chain kinase/MLCK) (mean ± SD; n = 10 cells per each perturbation). PM tension, plasma membrane tension-related pathway; ER/NE tension, endoplasmic reticulum/nuclear envelope tension-related pathway; [Ca^2+^]_out_, extracellular calcium entry; [Ca^2+^]_in_, release of calcium from intracellular sources. **B:** 3D images of the DAPI-stained nucleus (XZ views) from the same single live cell at 20, 10, and 5 μm confinement height, illustrating nuclear deformations. **C:** Left, DAPI-stained nucleus (XY view) from the same single live cell subjected to 10 μm confinement followed by height change to 5 μm, demonstrating nuclear expansion. Middle, measurements of projected nuclear surface area taken for the nucleus from the same single live cell at 10 *vs*. 5 μm confinement (n = 10 different cells; *p* value, paired t test). Right, measurements of nuclear volume taken from the same single live cell at 10 *vs*. 5 μm confinement (n = 10 different cells; *p* value, paired t test). Scale bar, 10 μm. **D:** The nuclear envelope (XY views) of the same single live cell at 20, 10, and 5 μm confinement height (scale bar, 5 μm. Zoomed images represent an example of local nuclear envelope unfolding from another cell. Scale bar, 2.5 μm), and quantifications of the degree of nuclear envelope folding measured as the excess of perimeter for the nuclear envelope (EOP_NE_) in the same single live cell confined to 10 and 5 μm (upper right, n = 10 different cells; *p* value, paired t test) or cells confined separately to 20, 10, and 5 μm (lower right, mean ± SD; n = 30 cells per each confinement height; *p* value, unpaired t test). **E:** The nuclear envelope (XY views) of the same single live cell confined to 5 μm and subsequently unconfined to 20 μm, and quantifications of the EOP_NE_ in the same single live cell confined to 5 μm and subsequently unconfined to 20 μm (n = 10 different cells; *p* value, paired t test). Scale bar, 5 μm. **F:** Representative nuclear envelope fluctuation curves obtained from cells at different confinement heights, and quantifications of the nuclear envelope fluctuation index in the same single live cell confined to 10 and 5 μm (n = 10 different cells; p value, paired t test) or cells confined separately to 20, 10, and 5 μm (mean ± SD; n = 30 cells per each confinement height; *p* value, unpaired t test). **G:** Nuclear pores on the nuclear envelope of the nucleus from the same single live cell at 10 and 5 μm confinement height, and quantifications of the inter-nuclear pore distance (NP-NP distance) measured in the same single live cell confined to 10 and 5 μm (n = 10 different cells; *p* value, paired t test). Scale bar, 0.5 μm. **H:** Subnuclear localization of the nuclear membrane tension sensor cPLA2 in the same single live cell at 10 and 5 μm confinement height, and quantifications of cPLA2’s stretch-sensitive accumulation at the nuclear envelope (ratio between the cPLA2 reporter signal at the nuclear envelope *vs*. the nucleoplasm (NE/NPM)) measured in the same single live cell confined to 10 and 5 μm (n = 10 different cells; *p* value, paired t test). Scale bar, 1.5 μm. **I:** A representative graph and time-lapse image sequence illustrating that nuclear envelope unfolding (accessed through measurements of EOP_NE_) temporally precedes the onset of active plasma membrane blebbing (measured as EOP_PM_) and associated with it contractile force response during the transition from 10 to 5 μm height. This phenomenon is observed in almost 100% of analyzed cases (n = 15 cells). Scale bar, 5 μm. **J:** Diagram illustrating that the degree of nuclear envelope folding (EOP_NE_) before cell confinement determines the height at which cells detect and respond to spatial confinement by generating contractile force (n = 20 cells). **K:** Proposed model: cells utilize the nucleus as an internal ruler for their height. Once cellular height is below a resting nuclear diameter, nucleus gets squeezed, which expands its envelope. When nuclear expansion reaches a critical level, nuclear envelope gets tensed (given that nuclear volume remains constant). This in turn stimulates nuclear stretch-sensitive proteins whose activity can initiate a signaling ultimately upregulating actomyosin contractility.

To understand how the nucleus and its envelope could contribute to triggering the sustained contractile response of cells to a specific confinement height, we characterized nuclear shape and deformation state at 20, 10 and 5 μm confinement (Fig. 2B). Based on chromatin staining, we observed that the nuclear volume remains relatively constant, while its projected surface area increased significantly between 10 and 5 μm confinement (Fig. 2C), suggesting a potential expansion of the nuclear envelope. To visualize the nuclear envelope more directly, we used cells expressing the inner nuclear membrane protein Lap2B fused to GFP. We observed that the nuclear envelope in rounded nonadherent cells (trypsinized cells assayed shortly after replating) displays large folds and wrinkles, which become less prominent at 10 μm and completely disappear once the cells are confined to 5 μm (Fig. 2D). To characterize this phenomenon quantitatively, we estimated the nuclear envelope folding index by measuring the excess of perimeter of the nuclear envelope (EOP_NE_) in the same single cell at various heights (n = 10 analyzed cells) or statistically comparing this parameter across populations of cells (n = 50 cells per population) confined to a specific height (Fig. 2D right graphs). Un-confining cells led to a rapid refolding of the envelope concomitant with the loss of the contractile response (Fig. 2E). These measurements suggest that within the range of confinement heights applied in our experiments, the nucleus maintains a constant volume by progressively unfolding its envelope until reaching a fully unfolded state at 5 μm. A higher degree of confinement and more severe nuclear compression results in a significant nuclear volume loss (with up to 50% of volume decrease at 3 μm (data not shown), also reported previously based on micropipette aspiration experiments (*24*)) and eventually nuclear envelope rupture events that become predominant below 3 μm confinement height (*25*).

As the nuclear envelope fully unfolds, it is likely to reach a state in which it stretches and becomes tensed. To estimate this parameter, we first measured the thermally and actively driven fluctuations of the nuclear envelope on the time scale of seconds (*26*). Using fast image acquisition mode for live cells expressing the Lap2B-GFP nuclear envelope marker and subpixel-resolution tracking of the nuclear envelope on consecutive images of the cells (see Methods), we found that the amplitude of fast fluctuations of the nuclear envelope systematically decrease as cells get more confined (Fig. 2F). This is consistent with an increase in the nuclear envelope tension. Imaging nuclear pores in the nuclear envelope of the same cell at various confinement heights showed that confinement from 10 to 5 μm causes neighboring nuclear pores to become more distant from each other, consistent with a stretching of the nuclear envelope that accommodates the pores (Fig. 2G). Finally, we observed that the nuclear-stretch activated phospholipase cPLA2, which senses lipid crowding in the nuclear envelope (*20*), remains in the nucleoplasm at 10 μm, and relocalizes to the nuclear envelope at 5 μm (Fig. 2H), a transition previously shown to be triggered by nuclear membrane tension increase and to correspond to cPLA2 enzyme activation (*20*). Overall, these observations suggest that confinement to 5 μm was inducing a stretching of the fully unfolded nuclear envelope.

To assess whether the height threshold at which cells display the active contractile response coincides with the induction of nuclear envelope stretching, we took advantage of the variety of nuclear shapes and folding states in the cell population and systematically investigated these parameters along with contractile force and cell morphology readouts. First, correlative recording of F-actin and the nuclear envelope (Fig. 2I bottom images) enabled us to observe that the nuclear envelope unfolding temporally precedes the onset of the contractile response manifested in vigorous cell blebbing (Fig. 2I top graph; Movie S3). This is compatible with a causal link between the nuclear envelope unfolding-stretching and the onset of a sustained contractile response. Second, we found that cells with more folded nuclei before confinement (larger EOP_NE_) start to contract at a lower confinement height (Fig. 2J), indicating that the degree of nuclear envelope folding sets the sensitivity threshold for the ability of cells to discriminate between different confinement heights.

Because we utilize cultured proliferating cells in our experiments, a source of cell-to-cell variability in responses to confinement could come from the cell cycle stage. A recent study showed that in cultured cycling cells, the nuclear envelope gets gradually smoother and tensed as the nucleus expands during the cell cycle progression from the G1 to the S/G2 phase (*26*). The cell cycle stage thus introduces a natural range of nuclear envelope states in a cell population. We used a FUCCI HeLa cell line with fluorescent cell cycle stage markers, confirming that G1 cells have a much higher excess of nuclear envelope perimeter than G2 cells (Fig. S5G). While the unconfined cell height and mechanical state (basal cortical tension and cell pressure) at 10 μm confinement were similar for rounded, nonadherent G1 and G2 cells (Fig. S5G), G2 cells required less confinement than G1 cells to trigger the contractile response (Fig. S5G bottom right graph). This result further confirms that the state and size of the nucleus defines a ruler to trigger the active contractile cell response. It also suggests that the nuclear ruler might render proliferating cells more or less sensitive to deformations depending on their cell cycle stage. This can provide a mechanistic basis for the recently described link between cell mechanics and cell cycle progression in multicellular epithelia and single cancer cells (*27, 28*), thus opening an exciting new avenue for future research.

Altogether, our results suggest the following working model (Fig. 2K): each single cell in the rounded state has a certain nuclear volume and an excess of nuclear envelope surface area, which promotes nuclear surface folding. When one of the dimensions of the cell is reduced below the resting nuclear diameter, the nucleus deforms and its envelope unfolds. Once the nuclear envelope reaches full unfolding, it stretches, potentially together with the perinuclear ER, leading to calcium release from internal stores and cPLA2 re-localization followed by the cPLA2 enzyme activation and production of the signaling lipid arachidonic acid/ARA. Both calcium ions and arachidonic acid are classical second messenger molecules with a well-known stimulatory effect on actomyosin contractility (*23, 29*–*31*), thus mechanistically linking the cell height and nuclear envelope stretching to cell contractility. Consistent with this model, the correlative live recording of cPLA2 localization, calcium (GCaMP6) and forces during changing the height from 10 to 5 μm, showed re-localization of cPLA2 within 20 seconds and intracellular calcium increase within less than a minute, both phenomena preceding the contractile response of the cell (Fig. S5H). We then decided to validate the nuclear ruler model by testing its direct predictions: the nuclear ruler pathway activation in cells squeezing through small holes, and its loss in enucleated cells as well as in cells with defects in nuclear envelope mechanics and morphology.

A first direct prediction of the nuclear ruler model is that as cells start squeezing through a tissue opening with a size smaller than a typical nuclear diameter, they should deform-unfold-and-stretch their nuclei as the nuclear shape gets less spherical, increasing the needed surface area for a given volume. This in turn would lead to an increase in cortical myosin II concentration when the nuclear ruler pathway turns on. To test this prediction in a controlled and quantitative manner, we flowed HeLa cells through confining microchannels with bottleneck constrictions. We found that nuclear deformations reflected in changes of the nuclear roundness index, indeed, precede myosin II accumulation at the cortex, and that this takes place when nuclear diameter is confined to approximately 5, but not 10 μm (Fig. 3A). To examine whether these dependencies can be observed in spontaneously migrating cells in a more physiological context, we looked at invasive HT1080 cells maneuvering through a 3D CDM. Plotting myosin II cortical accumulation against the plasma membrane excess of perimeter (i.e. cell blebbing and thus contractility measure), we verified that our measure of myosin II recruitment at the cell cortex is a good predictor of the degree of cell contractility in 3D (Fig. S6). We then found that this parameter scales with the extent of nuclear surface folding, which in turn is strongly dependent on nuclear height (Fig. 3B), very much similar to the results derived from our AFM- and microfluidics-based experiments. These data show that both imposed and spontaneous nuclear deformations observed in migrating cells correlate with the contractile response, suggesting that our working model could be applied to physiological contexts such as cells circulating in blood capillaries or cells migrating through dense tissues.

**Figure 3:**
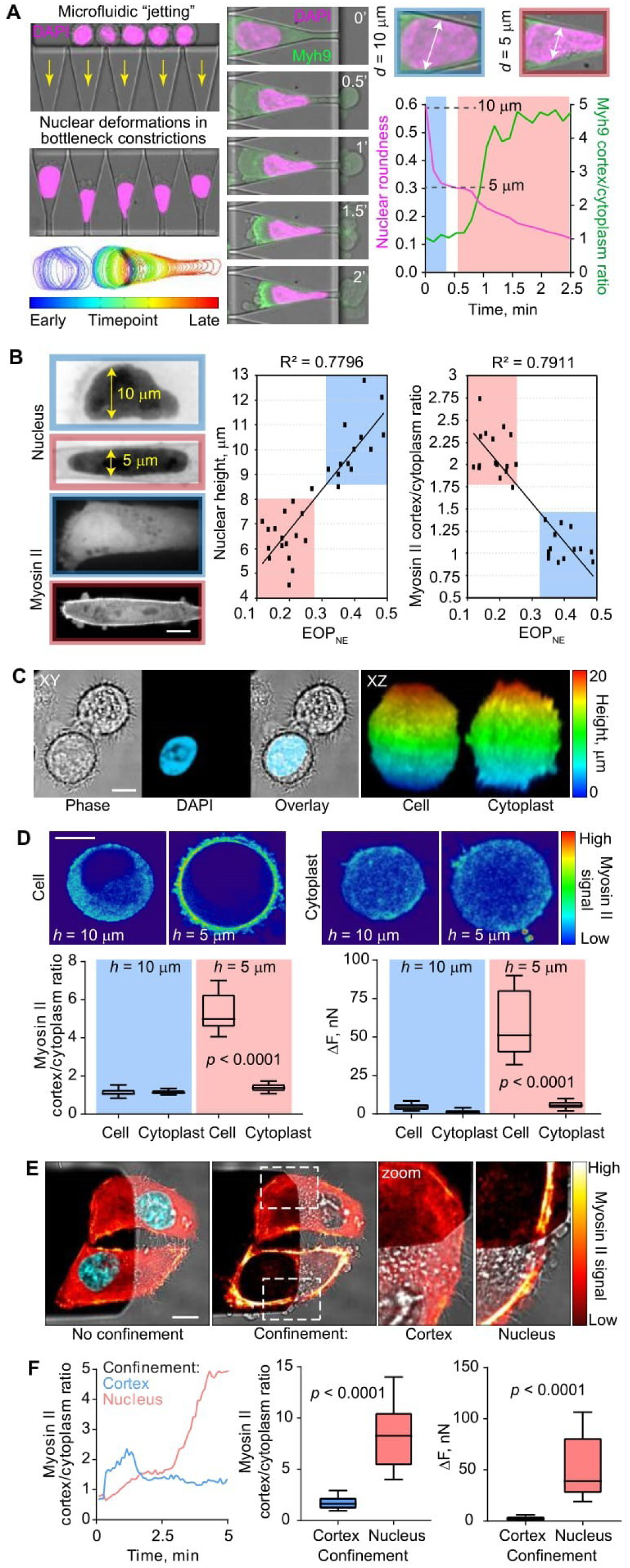
Cells “measure” their own height utilizing the nucleus as an internal ruler. **A:** Left, DAPI-stained HeLa-Kyoto cells microfluidically pushed through bottleneck PDMS constrictions to dynamically squeeze nuclei. Differentially-colored nuclear outlines demonstrate the evolution of nuclear deformation over time. Middle, visual correlation between various degrees of nuclear deformations and myosin II (Myh9) accumulation at the cortex at various time points. Right, a representative graph demonstrating that nuclear deformations (measured as nuclear roundness index) below a specific threshold (once nuclear width measures as 5 μm) induce cortical myosin II accumulation. This phenomenon is observed in ∼ 100% of analyzed cases (n = 30 cells). **B:** Left, representative images (XZ views, vertical cross-sections) of nuclei and myosin II in HT1080 cells expressing GFP-myosin light chain 2 and imaged within the 3D CDM. Middle, correlation between natural (as opposed to imposed in AFM or microfluidics experiments) nuclear height and the degree of nuclear envelope folding (EOP_NE_) in HT1080 cells imaged within the 3D CDM (n = 30 cells). Right, correlation between myosin accumulation at the cortex and the degree of nuclear envelope folding (EOP_NE_) in HT1080 cells imaged within the 3D CDM (n = 30 cells). Scale bar, 5 μm. **C:** Left, representative images (XY views) of nucleated cells and enucleated cytoplasts revealed by DAPI staining. Right, XZ views of nucleated cells and enucleated cytoplasts of a similar size selected for analyses. Scale bar, 5 μm. **D:** Quantifications of myosin II cortical accumulation and force response (ΔF) in nucleated cells (n = 10) and enucleated cytoplasts (n = 10) confined to 10 *vs*. 5 μm. Mean ± SD; *p* value, unpaired t test. Scale bar, 10 μm. **E:** Regioselective confinement of the nuclear region *vs*. the cortex of the cell lamella in spread cells. Scale bar, 10 μm. **F:** Quantifications of myosin II cortical accumulation and force response (ΔF) upon nuclear (n = 10) and lamellar cortex (n = 10) confinement in spread cells. Mean ± SD; *p* value, unpaired t test.

A second prediction of our working model is that removing the cell nucleus should affect the contractile response to confinement. We thus produced cytoplasts by cell enucleation using centrifugation (*32*). This resulted in a mixed population of enucleated cytoplasts and nucleated cells (Fig. 3C). Cytoplasts on average had a smaller volume compared to nucleated cells (Fig. S7A) but a rather similar height (note that a decrease in cell volume scales only to the cubic root of cell diameter). Thus, we were able to compare nucleated cells and enucleated cytoplasts of similar initial heights confined to 10 and 5 μm. While nucleated cells showed the expected contractile response at 5 μm, this was not the case for enucleated cytoplasts (Fig. 3D and Movie S4). Cytoplasts were not deficient in the contractile response pathway, because confining them further down to 1 μm triggered a strong contractile response (Fig. S7B). This is potentially due to direct compression and stretching of other endomembranes (e.g. endoplasmic reticulum (ER)) that remain present in enucleated cytoplasts (Fig. S7C). Endomembranes could take over the measuring function of the bulky nucleus at lower heights. To further demonstrate the difference in responsiveness to deformation provided by the nucleus/perinuclear ER *vs*. the cell cortex, we used spread cells and took advantage of the small size of the wedged cantilever tip to apply only a local deformation on the cell (Fig. 3E). This experiment confirmed that locally compressing the cell cortex in the lamellar region, even down to less than 1 micron (Fig. S8 and Movie S5), produces only a very transient response, while confining the part of the cell that contains the nucleus results in a rapid and sustained contractile response (Fig. 3F and Movie S6). Upon nuclear deformation, myosin II cortical recruitment occurred even in the region which was not directly confined (Fig. 3F). This showed that the contractile response is not due to the deformation *per se*, but rather due to a signal which could get released locally and propagate away from the nucleus (indeed, the increase in contractility in the non-confined part manifested with a slight delay compared to the nuclear region, Movie S6). In conclusion, consistent with our working model, the nucleus is required to set the size at which the contractile response is triggered and the deformation of nuclear or nucleus-associated compartments is necessary to trigger the sustained response.

We further tested the nuclear ruler model by more directly affecting two key parameters: the stiffness and the folded state of the nuclear envelope. As expected, Lamin A/C depleted cells (Fig. S9A left western blot) contained more deformed nuclei with a more floppy envelope (Fig. 4A). The depleted cells also displayed a high rate of nuclear envelope ruptures at 5 μm confinement (Fig. 4A bottom left graph), a phenomenon which could on its own relax tension in the envelope. Consistently, unlike in control cells (Fig. 2F), fluctuations of the nuclear envelope in Lamin A/C-depleted cells did not decrease in response to 20-10-5 μm confinement, suggesting that the envelope remained floppy at all confinement heights (Fig. 4A top right graph). Confinement of depleted cells to 5 μm did not trigger intracellular calcium release nor did it increase levels of ARA production normally expected from a tensed nuclear/perinuclear ER membrane compartment (Fig. S9A middle graphs). Consistently, while basal mechanics (cortical tension and intracellular pressure) of the depleted cells were unaltered at 10 μm (Fig. S9A right graphs), confinement of these cells to 5 μm was not accompanied by a contractile force response (Fig. 4A bottom right graph). This is again consistent with our working model for the nuclear ruler. It further suggests that the level of Lamin A/C in the nuclear lamina, which varies to a great extent in different cell types and environmental conditions, can modulate the response of cells to spatial confinement by affecting the mechanical properties of the nuclear envelope.

**Figure 4:**
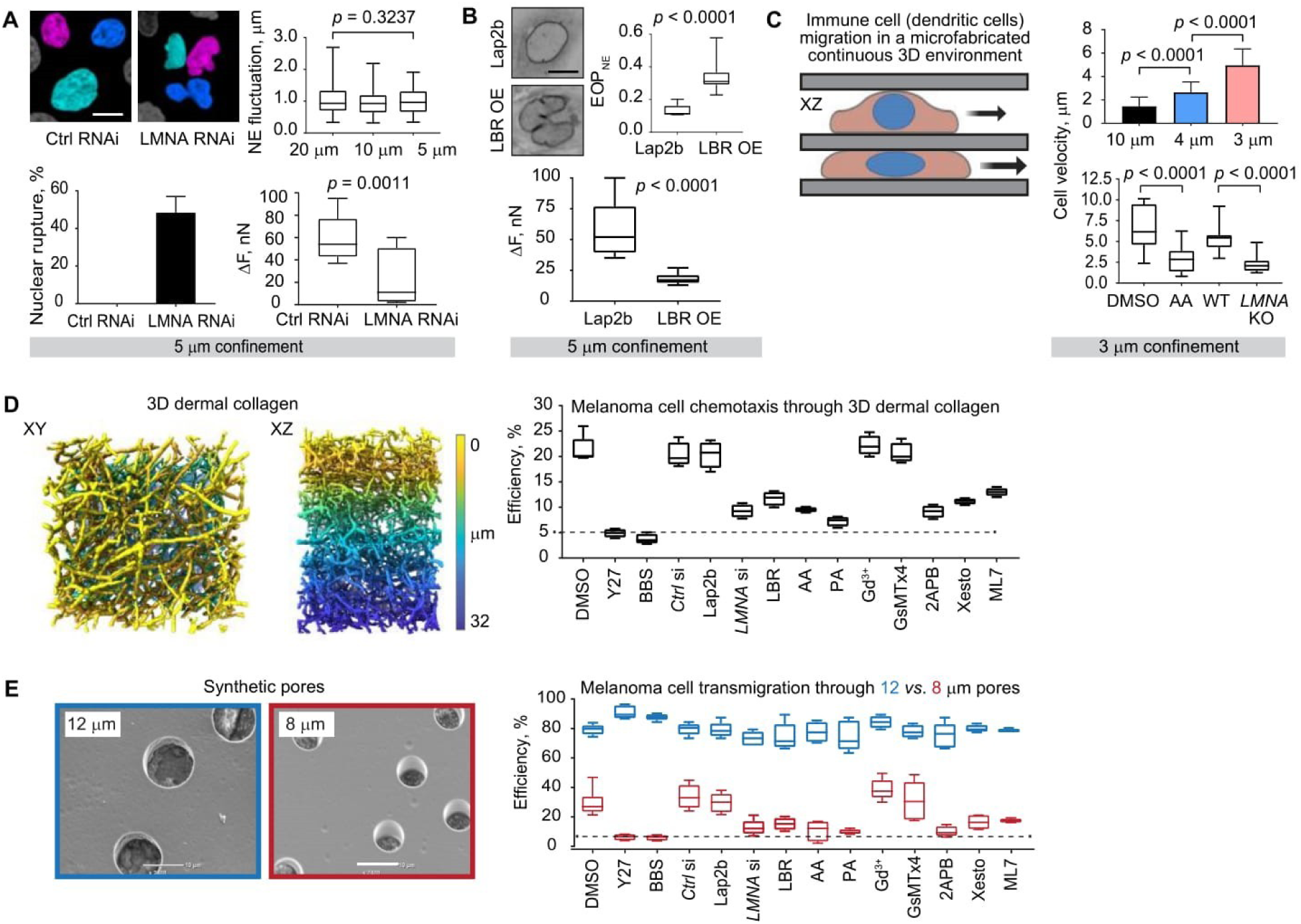
Testing the limits of the nuclear ruler and its physiological role during cell motility in complex environments. **A:** Top left, representative images of DAPI-stained nuclei of HeLa-Kyoto cells in a Petri dish treated with control and LMNA siRNA. Top right, quantifications of nuclear envelope fluctuation index in LMNA siRNA-treated cells under 20, 10, and 5 μm confinement (mean ± SD; n = 20 cells per each height; *p* value, unpaired t test). Bottom left, percentage of control siRNA-treated cells (n = 15) and LMNA siRNA-treated cells (n = 15) displaying nuclear rupture under 5 μm confinement. Bottom right, force response (ΔF) to 5 μm confinement measured in control siRNA-(n = 10) and LMNA siRNA-treated cells (n = 15). Mean ± SD; *p* value, unpaired t test. Scale bar, 10 μm. **B:** Top, representative images of nuclei in control cells expressing Lap2b-GFP *vs*. overexpression (OE) of LBR-GFP in a Petri dish, and measurements of the degree of nuclear folding (EOP_NE_) in these cells (mean ± SD; n = 10 cells per each case; *p* value, unpaired t test). Bottom, force response (ΔF) to 5 μm confinement measured in Lap2b-GFP (n = 10) and LBR-GFP OE cells (mean ± SD; n = 15; *p* value, unpaired t test). Scale bar, 10 μm. **C:** Primary immature mouse bone marrow-derived dendritic cells acquire high-speed, ballistic-like motion once spatially confined to 3 (n = 35 cells) but not 4 (n = 35 cells) or 10 (n = 20 cells) μm between two parallel surfaces mimicking interstitial tissue spaces. The gain in cell velocity observed at 3 μm height in control cells (DMSO and wild-type (WT)) is cancelled upon inhibition of the nuclear envelope tension sensor cPLA2 with AA (AACOCF3) or nuclear LMNA knock-out (KO). Mean ± SD; n = 20 cells per each experimental condition; *p* value, unpaired t test. **D:** 3D super-resolution microscopy images of dermal collagen lattices displaying pores and constrictions of different sizes, and quantifications of the percentage of human melanoma cells A375 able to chemotactically transmigrate through the lattice (1 mm thick) in a proteolysis-independent manner (determined by the presence of the matrix metalloproteinase (MMP) inhibitor GM6001) in the following conditions: control (DMSO, ctrl siRNA, and Lap2b expression), basal actomyosin contractility inhibition with Y27632 and blebbistatin (Y27 and BBS), nuclear LMNA depletion (LMNA si), nuclear LBR-GFP overexpression (LBR), nuclear envelope tension sensor cPLA2 inhibition with AACOCF3 or an alternative inhibitor PACOCF3 (AA and PA), inhibition of stretch-sensitive ion channels on the plasma membrane with GdCl_3_ or an alternative inhibitor the peptide GsMTx4 from the tarantula venom (Gd^3+^ and GsMTx4), inhibition of the stretch sensitive Ca^2+^-channels InsP3R present on the perinuclear ER and nuclear membranes with 2APB or an alternative inhibitor Xestospongin C (2APB and Xesto), and Ca^2+^-calmodulin-dependent myosin light chain kinase/MLCK inhibition with ML7 (ML7). **E:** Scanning electron microscopy images of polycarbonate membranes with 12 and 8 μm pores in diameter (scale bar, 10 μm), and quantifications of the percentage of human melanoma cells A375 able to chemotactically transmigrate through the differentially-sized pores in the conditions replicating those specified in **D**.

The inner nuclear membrane and ER membrane protein lamin B receptor/LBR is known to control nuclear envelope folding in a dose-dependent manner (*33*). Its overexpression leads to a massive perinuclear ER expansion and overproduction of nuclear envelope membranes (*34*–*36*). This in turn provides additional nuclear envelope surface area to accommodate excess membrane protein, as evidenced from our measurements of EOP_NE_ for Lap2b-GFP (control)-expressing cells *vs*. LBR-GFP overexpressing cells (Fig. 4B images and top graph). This parameter remained high in LBR-GFP overexpressing cells upon 10-to-5 μm confinement (Fig. S9B images and top graphs). This shows that the nuclear envelope did not fully unfold at 5 μm in LBR-GFP overexpressing cells. Consistently, LBR overexpression abrogated both cytoplasmic calcium increase and levels of ARA production at 5 μm (Fig. S9B bottom left graphs). While LBR-GFP overexpressing cells displayed basal cortical mechanics similar to control cells at 10 μm confinement (Fig. S9B bottom right graphs), the cells had significantly impaired contractile force response at 5 μm (Fig. 4B bottom graph). Overall, these experiments demonstrate that nuclear envelope unfolding triggers the contractile response upon confinement. Nuclear envelope folding and unfolding thus constitutes a key element of the nuclear ruler mechanism. It further suggests that modulation of nuclear envelope components that affect the extent of nuclear envelope folding (e.g. LBR and SUN2 (*37*)) could allow different cell types to trigger responses to various levels of confinement, or to measure different ranges of sizes, depending on their function.

Finally, we wanted to assess the functional relevance of the nuclear ruler. Mutations in the *LBR* gene lead to autosomal genetic disorders such as Pelger-Huët anomaly and Greenberg dysplasia (*38*). Immune cells from human patients carrying an *LBR* heterozygous dominant mutation display nuclei with abnormal shapes and inability to squeeze through small, but not large confines during *in vivo* chemotactic migration previously assessed in clinical studies through the skin-window technique (*39*). Based on our data, we propose that LBR-mediated changes in the nuclear ruler function could result in dysregulation of signals from the nuclear envelope to the actomyosin cytoskeleton and impede confined but not basal cell migration. Our conjecture is that cells migrating in complex 3D environments and encountering various tissue confines could utilize the nuclear ruler to discriminate between small and large openings (*3*). If the opening is smaller than the nuclear size, nuclear deformation and stretching can trigger actomyosin contractility providing a motive force for the cell to squeeze through the opening to avoid entrapment, suggesting a function for the nuclear ruler in cell migration.

Generalizing this concept, we hypothesized that the nuclear ruler mechanism can be used by migrating cells to increase their propulsion force when space becomes limited, explaining the switch to fast amoeboid migration upon confinement of slow mesenchymal cells, which we previously reported (*10*). To test if this can also apply to professionally migrating cells such as primary immune cells, we confined immature mice bone marrow-derived dendritic cells at various heights using our 2D cell confiner device (*40*). We observed the predicted increase in migration speed upon confinement (Fig. 4C). This speed increase was lost upon inhibition of cPLA2 (AA) or depletion of Lamin A/C (*LMNA* KO, Fig. 4C bottom graph), suggesting that immature dendritic cells sense spatial confinement and adapt their contractility relying on the nuclear ruler pathway.

To further test the function of the nuclear ruler mechanism in cell migration, we assessed the protease-independent ability of human skin melanoma cells, a well-established cellular model employing contractility-driven amoeboid motion *in vivo* (*41*), to chemotactically transmigrate (Fig. S10A) through 3D dermal collagen gels. The 3D collagen lattices (Fig. 4D left images) displayed pore sizes ranging from 1 to 12 μm (Fig. S10B), thus representing a heterogeneous, mechanically restrictive environment in which cells are expected to deform their nuclei. Consistent with our experimental confinement data, perturbations specific to the nuclear envelope/perinuclear ER membrane tension and calcium-dependent MLCK, as well as biophysical properties of the nuclear envelope, affected the efficiency of melanoma chemotactic transmigration through 3D collagen (Fig. 4D right graph). To more directly test whether this outcome was determined by the nuclear ruler and restrictive size of pores in the heterogeneous 3D collagen lattice, we compared transmigration efficiency of melanoma cells through synthetic polycarbonate membranes with nonrestrictive (d = 12 μm) *vs*. restrictive (d = 8 μm) pores (Fig. 4E left images and Fig. S10C). We found that melanoma cells rely on basal actomyosin contractility or pathways associated with the nuclear envelope/perinuclear ER membrane tension and calcium-dependent MLCK as well as biophysical properties of the nuclear envelope only when transmigrating through 8-μm pores (Fig. 4E right graphs) that are smaller than the average size of the nucleus in melanoma cells (diameter of 11 ± 2 μm, n = 100 cells). These results suggest that thanks to the nuclear ruler, migratory cells can utilize the energetically costly actomyosin contractility motor on demand, when local cell environment becomes excessively restrictive to cell migration.

Collectively, our data establish a fundamentally new, nongenetic function for the nucleus as an internal ruler. Relying on this ruler, cells can measure the degree of their environmental confinement and rapidly tailor specific behaviors to adapt to the confinement at timescales shorter than changes in gene expression. In the context of cell migration, such tailored cellular behaviors might help cells to avoid environmental entrapment, which is relevant to cancer cell invasion, immune cell patrolling of peripheral tissues, and progenitor cell motility within a highly crowded cell mass of a developing embryo. The nuclear ruler mechanism defines an active function for the nucleus in cell migration, potentially explaining why enucleated cells show a poor motile capacity in dense collagen gels (*42*). Establishing the nucleus as an internal ruler of extracellular environment opens up new exciting avenues of research not only in the field of 3D cell migration, but also tissue homeostasis and developmental biology. Tissue cells such as epithelial cells might utilize the nucleus to sense local cell density, which is critically involved in tissue formation, homeostasis, turnover, and wound healing.

## Acknowledgments

The authors wish to acknowledge I. Poser and A. Hyman (Max Planck Institute of Molecular Cell Biology and Genetics, Dresden, Germany) for providing stable BAC transgenic HeLa cell lines expressing various fluorescent protein markers, as well as P. Niethammer (Sloan-Kettering Institute, New York, New York, USA) for sharing the HeLa cell line that stably expresses cPLA2α-mKate2, and V. Sanz-Moreno (Queen Mary University of London, London, UK) for providing A375P cells. LBR pEGFP-N2 (646) was a gift from E. Schirmer (Addgene plasmid #61996; http://n2t.net/addgene:61996; RRID: Addgene_61996). pGP-CMV-GCaMP6s was a gift from D. Kim & GENIE Project (Addgene plasmid # 40753; http://n2t.net/addgene:40753; RRID: Addgene_40753). pBOB-EF1-FastFUCCI-Puro was a gift from K. Brindle & D. Jodrell (Addgene plasmid #86849; http://n2t.net/addgene:86849; RRID: Addgene_86849). We are grateful to B. Baum (University College London, London, UK) and D. Gerlich (Institute of Molecular Biotechnology of the Austrian Academy of Sciences, Vienna, Austria) for comments on the manuscript. We also thank G. Charras (University College London, London, UK) for the advice to estimate inter-nuclear pore distance in nuclei at various degrees of spatial confinement, and N. Carpi (Institut Curie, Paris, France) for excellent technical assistance with experiments. Some illustrations accompanying figures of the present manuscript were partially generated using the BioRender.com online tool;

## Funding

The research leading to these results has received funding from the People Programme (Marie Curie Actions) of the European Union’s Seventh Framework Programme (FP7/2007-2013) under REA grant agreement n. PCOFUND-GA-2013-609102, through the PRESTIGE programme coordinated by Campus France. A.J.L. was supported by the Marie Curie & PRESTIGE Fellowship (grant 609102), London Law Trust Medal Fellowship (grant MGS9403), and a Career Grant for Incoming International Talent (grant 875764) from the Austrian Research Promotion Agency (FFG). I.Z. was supported by a Metchnikov Fellowship from the Franco-Russian Scientific Cooperation Program;

## Author contributions

A.J.L., C.J.C., D.J.M. and M.P. designed the project. A.J.L. and C.C. performed all key experiments and analyzed the data. D.C. analyzed most of force spectroscopy data and developed analytical approaches to estimate the degree of nuclear membrane folding. Z.A. and A.M.L-D. performed experiments with primary mouse immature dendritic cells. M.M. performed experiments with A375P cells. G.N. performed experiments with HeLa-cPLA2α cells. N.S. performed experiments with HeLa-Lap2B cells and developed analytical approaches to estimate the degree of nuclear membrane fluctuation. J.M.G-A. and I.Z. performed western blot analysis of LMNA knockdown efficiency as well as experiments on the effect of transcription and translation inhibition in HeLa cells. J.M.G-A. additionally assisted with manuscript preparation for submission. A.B. and N.M. performed shRNA-mediated LMNA knockdown in HeLa cells. M.K.D., E.S.W., and R.F. analyzed 3D collagen data. R.J.P. performed experiments with HT1080 cells in 3D CDMs. A.J.L., C.C., D.J.M. and M.P. wrote the manuscript. All authors discussed the results and implications, and commented on the manuscript at all stages;

## Competing interests

The authors declare no competing financial interests;

## Data and materials availability

All data is available in the main text or the supplementary materials.

