## Supplementary Materials for "The nucleus acts as a ruler tailoring cell responses to spatial constraints"

**This PDF file includes:**

Materials and Methods
Figs. S1 to S10
Tables S1
Captions for Movies S1 to S6

**Other Supplementary Materials for this manuscript include the following:**

Movies S1 to S6

### Materials and Methods

Cell Culture

Human cervical adenocarcinoma cells HeLa-Kyoto stably expressing myosin IIA (Myh9)-GFP and LifeAct-mCherry or Myh9-GFP and the plasma membrane-targeting CAAX box fused to mCherry, or Lap2b-GFP, Nup107-GFP, HeLa (CCL-2) cells stably expressing cPla2-mKate2, human fibrosarcoma cells HT1080 expressing GFP-myosin light chain 2, primary human foreskin fibroblasts (HFFs), human melanoma cells A375P, *N-rasV12* oncogene-transformed rat liver epithelial cells IAR-2, and canine kidney epithelial cells MDCK-2 were maintained in DMEM/F12 supplemented with 10% FBS (Invitrogen) at 37°C and 5% CO2. Human epidermal stem cells (HESCs) were cultured as previously described (*1*). All cell lines were tested for mycoplasma contamination using MycoScope™ PCR Mycoplasma Detection Kit (Genlantis). Mouse bone-marrow derived dendritic cells (DCs) were obtained by culturing bone marrow cells (from both male and female control or LMNA KO mice) for 10-11 days in complete DC medium (IMDM medium supplemented with fetal calf serum (FCS, 10%), glutamine (20 mM), penicillin-streptomycin (100 U/mL), ß-mercaptoethanol (50 µM) and granulocyte-macrophage colony-stimulating factor (50 ng/mL)-containing supernatant obtained from transfected J558 cells.

Transfection procedure, expression vectors, and siRNA oligonucleotides

Cells were transfected with plasmid DNA using Lipofectamine™ LTX reagent (Invitrogen) transiently or stably, according to manufacturer's protocol. For RNA interference experiments, cells were transfected with siRNA oligonucleotides using Lipofectamine® RNAiMAX reagent (Invitrogen), according to manufacturer's protocol.

The following expression vectors were used for plasmid DNA transfections: empty vector pEGFP-C1 (Clontech); Addgene plasmids: 61996 LBR pEGFP-N2 (646) (*2*), 40753 pGP-CMV-GCaMP6s (*3*), 86849 pBOB-EF1-FastFUCCI-Puro (*4*).

To knockdown LMNA/C, cells were transfected with nontargeting siRNA (control) or validated ON-TARGETplus SMARTpool siRNA reagents (Dharmacon) targeting human-specific LMNA mRNA (cat. # L-004978-00-0005). Cells were analyzed 72 h post-transfection using standard Western blot or immunofluorescent analysis protocols. Based on quantitative densitometry of proteins, the knockdown efficiency was estimated as 84.7 ± 2.5% (3 repeats). Additionally, a lentiviral vector plasmid encoding a nontargeting shRNA (control) or shRNA targeting human-specific LMNA mRNA designed in house and standard lentivirus-based transduction-infection protocols were used to achieve LMNA/C knockdown in HeLa cells.

Drug treatments

The following pharmacological inhibitors and chemical compounds were used: 10 µM ROCK-mediated contractility inhibitor Y-27632 (Y27) (EMD), 10 µM myosin II ATPase inhibitor blebbistatin (BBS) (Toronto Research Chemicals), 20 µM $Ca^{2+}$-sensitive myosin light chain kinase/MLCK inhibitor ML-7 (Sigma-Aldrich), 1 mM apoptosis inducer hydrogen peroxide ($H_2O_2$), 1 µM transcription inhibitor triptolide (TRP) (Tocris Bioscience), 50 µg $ml^{-1}$ translation inhibitor cycloheximide (CHX) (Sigma-Aldrich), 10 µM nonspecific plasma membrane permeability marker propidium iodide (PI) (Sigma-Aldrich), 0.4 mM plasma membrane tension reducer sodium deoxycholate (DOCL) (Sigma-Aldrich), 10 µM gadolinium (III) chloride ($Gd^{3+}$) or 5 µM peptide GsMTx4 from the tarantula venom affecting mechanosensitive ion channels on the plasma membrane (Tocris Bioscience), 2 mM extracellular

$Ca^{2+}$ chelator BAPTA (Sigma-Aldrich), 10 µM intracellular $Ca^{2+}$ chelator BAPTA-AM (Sigma-Aldrich), 10 µM ionomycin (IOM) directly facilitating the transport of $Ca^{2+}$ across the plasma membrane (Sigma-Aldrich), 70 µM signalling lipid arachidonic acid (ARA) (a product of enzymatic activity of the nuclear envelope stretch-sensitive enzyme cPLA2) activating actomyosin contractility (Cayman Chemical), 20 µM AACOCF3 (AA) or 10 µM PACOCF3 (PA) inhibiting the nuclear envelope stretch-sensitive enzyme cPLA2 (Tocris Bioscience), 100 µM 2APB or 10 µM Xestospongin C (Xesto) blocking stretch-activated inositol triphosphate receptors (InsP3Rs) on the ER/nuclear membranes (Tocris Bioscience), and 20 µM broad-spectrum matrix metalloproteinase inhibitor GM6001 (Merck Millipore). Growth medium was supplemented with 1% DMSO (vol/vol) (Sigma-Aldrich) in control experiments.

Western blotting

Cells were collected and resuspended in Laemmli buffer. Proteins were separated using sodium dodecyl sulfate polyacrylamide gel electrophoresis (SDS–PAGE) and transferred onto PVDF membranes. After incubation with primary (Lamin A/C Antibody #2032 (Cell Signaling Technology) and GAPDH antibody #ab9483 (Abcam)) and secondary (IRDye® and VRDye™ (LI-COR)) antibodies, the membranes were visualized using Odyssey® CLx Infrared Imaging System (LI-COR).

Single-cell flat AFM-based confinement coupled to live cell imaging

Trypsinized cells were resuspended in CO2-independent, phenol red-free DMEM/F-12 medium supplemented with 10% FBS (Invitrogen) and plated on glass-bottomed 35-mm dishes (Fluordishes). Experiments with non-adherent cells were initiated 30 minutes after cell plating to allow for cell sedimentation. Spread cells were obtained 6 hours post cell plating. Dishes with cells were mounted in a dish heater (JPK Instruments) and kept at 37 °C under an inverted light microscope (Axio Observer.Z1; Zeiss) equipped with a confocal microscope unit (LSM 700; Zeiss) and the atomic force microscope (AFM) head (CellHesion 200; JPK Instruments).

Focused ion beam (FIB)-sculpted, flat silicon microcantilevers were processed and calibrated as described in (*5*). The microcantilevers were fixed on a standard JPK glass block and mounted in the AFM head. The cantilever was lowered on the cell to a preset height with a constant speed of 0.5 $\mu m \cdot s^{-1}$, and the resulting varying force and cantilever height were recorded over time. At the same time, differential interference contrast (DIC) and fluorescence images at the midplane of the confined cell were recorded every 5 seconds using a 63× water immersion objective. All microscopy equipment was placed and experiments were carried out in a custom-made isolation box at 37 °C (The Cube; Life Imaging Services).

Determination of cell pressure and cortical tension

Cell geometry, pressure, and cortical tension were measured based on AFM and imaging data as described in (*5*) and (*6*).

Production of cytoplasts

Enucleated cells were generated as described in (*7*) and (*8*).

Microfabrication-based confinement of cell populations

To obtain large quantities of confined cells for cell population or biochemical studies, cell confinement was performed using a home-made device (*9*) consisting of a suction cup made in polydimethylsiloxane (PDMS, RTV615, GE) used to press a confining coverslip bearing PDMS microspacers (micropillars) on top of the culture substrate populated with cells. The height of the micropillars (10 µm *vs.* 5 µm) determines the height for spatial confinement of cells between the coverslip and the substrate. A version of the cell confiner adapted to multi-well plates was used to perform multiple experiments in parallel (*10*). The molds for the PDMS micro-spacers were fabricated following standard photolithography procedures. The surface of the confining side was always treated with non-adhesive pLL-PEG (SuSoS).

<u>Assaying activation of apoptosis in live cells</u>

To detect levels of active apoptotic caspases, the Image-iT LIVE Red Poly Caspases detection kit based on a fluorescent inhibitor of caspases (FLICA) methodology (I35101, Molecular probes) was used according to the manufacturer's protocol.

<u>Biochemical measurements of arachidonic acid (ARA) release</u>

Cells were confined using microfabricated devices as described in the subsection “Microfabrication-based confinement of cell populations”. Confinement was released and the cells were immediately extracted with Dole's solution (heptane, isopropyl alcohol, 1 N sulfuric acid; 10:40:1). Pentafluorobenzyl esters of the fatty acids were prepared and quantified by gas chromatography-mass spectometry with reference to an internal standard of d8-AA as described in (*11*).

<u>Epithelial monolayer stretching</u>

A custom-made stretching device was used to perform epithelial monolayer stretching experiments as described in (*12*).

<u>Chemotactic transmigration assays</u>

Serum-starved A375P cells were harvested and transferred in serum-free medium to the upper compartment of 5, 8 (Cat. # 3421 and 3428, Corning), or 12 (Cat. # CBA-107, Cell Biolabs Inc.) µm-pore polycarbonate membrane inserts (transwells). Cell density was adjusted according to the specific area of each transwell with 5×104, 1.5×105, and 7.5×105 cells added to 5, 12, and 8 µm transwells, respectively. Cells were allowed to transmigrate toward the lower compartment containing 10% FBS for 12 h. Transmigration efficiency was calculated as number of cells at the lower compartment divided by the number of cells added to the upper compartment of a transwell.

To examine the ability of A375P cells to chemotax through 3D collagen gels, atelopeptide fibrillar bovine dermal collagen (Cat. # 5005-B; PureCol, Advanced BioMatrix) was prepared at 1.7 mg $ml^{-1}$ in DMEM and allowed to polymerize in the upper compartment of the 12 µm-pore polycarbonate membrane insert (Cat. # CBA-107, Cell Biolabs Inc.). Serum-starved cells were seeded on top of the collagen pad in serum-free medium, allowed to adhere and transmigrate through the collagen layer and the membrane toward the lower compartment containing 10% FBS for 24 h. Transmigration efficiency was calculated as number of cells at the lower compartment divided by the number of cells added to the upper compartment of the transwell.

<u>Generation of 3D cell-derived matrices (CDMs)</u>

HFFs were plated at high density on gelatin-coated and glutaraldehyde-treated 35 mm (4 x $10^5$ cells, MatTek) or 50 mm (5.7 x $10^5$ cells, Warner Instruments) glass-bottom dishes. Cultures were maintained for 10 days, adding new media with 50 µg/ml ascorbic acid every other day. The matrices were denuded of cells by adding extraction buffer (20 mM $NH_4OH$ and 0.5% Triton X100 in PBS) for 10 min at room temperature and washed with PBS.

Quantitative image analysis

Fluorescence intensity levels (mean gray values, a.u.) were obtained from background subtracted images using the ImageJ/Fiji software (NIH, http://rsb.info.nih.gov/ij/index.html).

The Imaris (version 8.3) image analysis software (Bitplane) was utilized to measure nuclear volume reconstructed from 3D confocal microscopy images.

The dimensionless parameter excess of perimeter (EOP) was calculated to estimate the amount of nuclear envelope (NE) or plasma membrane (PM) area stored in macro- and micro-folds. To calculate EOP, we first obtained values for perimeter (P) and surface area (A) from 2D images taken at the maximum radius of the nucleus (NE marker Lap2b-GFP labelling)/cell (PM marker CAAX box-mCherry labelling). Next, we introduced $R_0$ as the radius of the circle defined by the area A, which allowed us to compute EOP as the ratio between P and $2\pi R_0$. EOP values of a highly folded object tend to be close to 1, while EOP of an object with a smooth surface tend to be close to 0.

To measure NE fluctuations, nuclei of live cells expressing Lap2b-GFP were recorded using a high frame rate acquisition mode (250 msec/frame) for 5 minutes. The position of each nucleus was corrected for linear and rotational drift using the Stackreg plugin of the ImageJ/Fiji software. Upon drift correction, the edge of the nucleus was registered at a given angle. To estimate the amplitude of NE fluctuations over time, standard deviation of the edge from its mean position was calculated and considered as total fluctuation of the NE. The edge position was tracked at different angles for each nucleus to obtain mean NE fluctuation values expressed in µm. The same approach can be applied to measure PM fluctuations. A floppy, and thus less tensed, membrane is expected to fluctuate more, while a membrane under tension exhibits no or significantly diminished fluctuations.

The distance between neighboring nuclear pores (NP-NP distance) was estimated for the same living cell expressing a NP maker Nup107-GFP at different degrees of spatial confinement. To this end, a medial confocal slice of the nucleus was obtained to then trace individual NPs along the NE using the Linescan function of the MetaMorph (version 7.7) software (Molecular Devices). Each individual NP was resolved as a local maximum of fluorescence intensity on the Linescan diagram. The distance between nearest local maxima was represented as NP-NP distance.

Cell speed was analyzed using the manual tracking plugin of ImageJ/Fiji.

To measure collagen pore sizes, collagen fibers were imaged *via* axially swept light-sheet microscopy (*13*). The fibers were then detected from the deconvolved images by applying a steerable filter (*14*) followed by non-maximum-suppression. We then calculated pore sizes using a custom-written Matlab code implementing the algorithm described in (*15*).

Statistics and reproducibility of experiments

Unless stated otherwise, statistical significance was determined by two-tailed unpaired or paired Student's t-test after confirming that the data met appropriate assumptions (normality, homogenous variance and independent sampling). Statistical data are presented as average ± either SEM or SD. Sample size (n) and p-value are specified in the text of the paper or figure legends. Samples in most cases were defined as the number of cells counted/examined within multiple different fields of view on the same dish/slide, and thus represent data from a single sample within a single experiment. When data from a single sample are shown, they are representative of at least three additional samples from independent experiments.

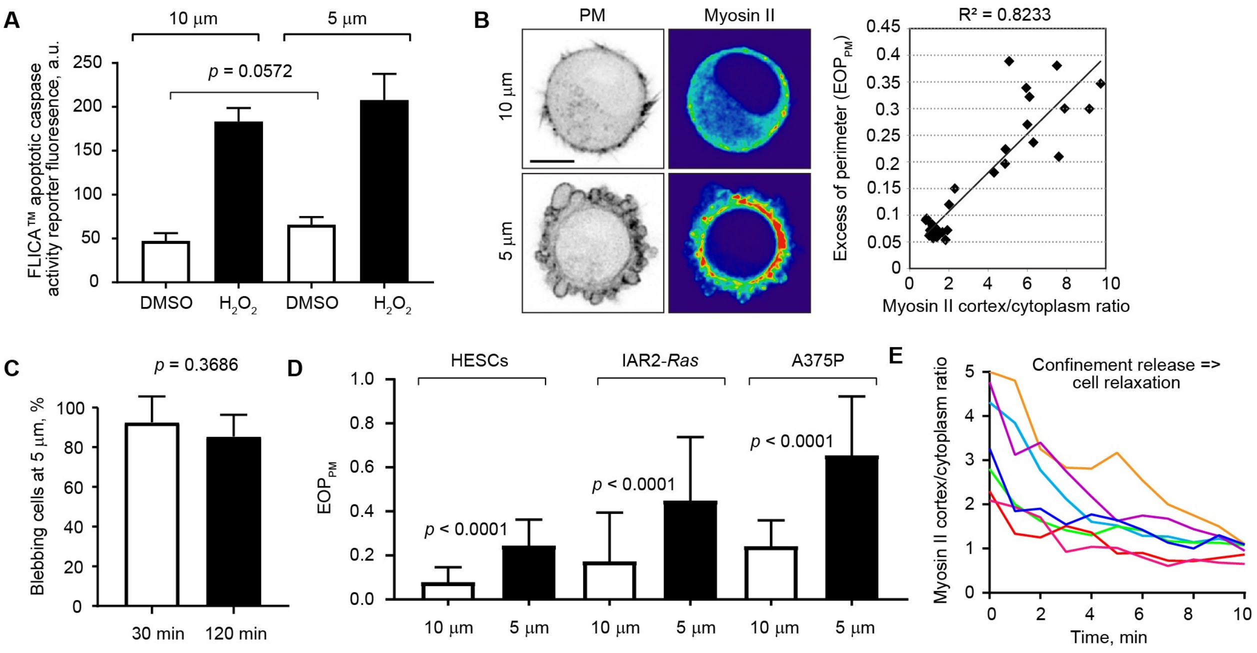


**Fig. S1.**

**Non-apoptotic plasma membrane blebbing associated with the contractile response to threshold confinement is sustained over time, observed in different cell types, and represents a reversible phenomenon.**

**A:** Comparing apoptosis induction efficiency (quantified as average fluorescence intensity of the FLICA apoptotic caspase activity reporter in cells) in response to spatial confinement of cells to 10 and 5 μm *vs.* treatment of cells with a classical apoptosis inducer hydrogen peroxide as a positive control. Mean ± SD; n = 85 cells from 3 independent fields of view per each experimental condition; *p* value, unpaired t test.

**B:** The plasma membrane (PM) blebbing index, measured as the excess of plasma membrane perimeter ($EOP_{PM}$), correlates with cortical accumulation of myosin II in cells under 5 μm confinement (n = 35 cells). Scale bar, 10 μm.

**C:** Percentage of blebbing cells 30 and 120 minutes post 5 μm-confinement. Mean ± SD; n = 120 cells per each time point from 3 independent fields of view; *p* value, unpaired t test.

**D:** Quantifications of the plasma membrane (PM) blebbing index measured as the excess of plasma membrane perimeter ($EOP_{PM}$) at 10 and 5 μm confinement in primary human epidermal stem cells (HESCs, n = 45 cells per each height), immortalized *N-ras*-transformed rat liver hepatocytes IAR-2 (n = 150 cells per each height), and human melanoma cell line A375P (n = 40 cells per each height). Mean ± SD; *p* value, unpaired t test.

**E:** Quantifications of myosin accumulation at the cortex in response to release of spatial confinement (n = 7 cells).

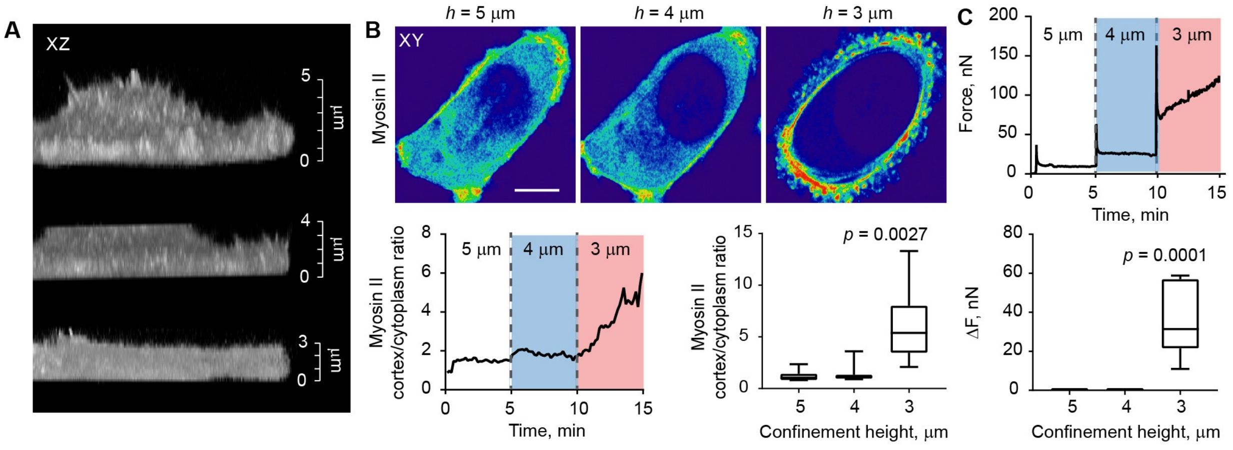


**Fig. S2.**

**Cell adhered to the substrate sense spatial confinement, but due to spreading-associated flattening respond to critical confinement at heights lower than those established for rounded nonadherent cells.**

**A:** 3D images (XZ views) of the same single live cell at 5, 4, and 3 µm confinement height.

**B:** Top, myosin II fluorescence intensity levels in the same single live cell (XY views) sequentially subjected to 5, 4, and 3 µm confinement. Bottom left, a representative graph illustrating dynamic relocalization of myosin II from the cytoplasm to the cell cortex in the same single live cell sequentially confined to 5, 4, and 3 µm. Bottom right, statistical analysis of cortical accumulation of myosin II detected in different cells at 5, 4, and 3 µm confinement. Mean ± SD; n = 7 cells per each height; *p* value, unpaired t test. Scale bar, 10 µm.

**C:** Top, a representative force response curve of the same single live cell sequentially confined to 5, 4, and 3 µm. Bottom, statistical analysis of force response output (ΔF) detected in different cells at 5, 4, and 3 µm confinement. Mean ± SD; n = 10 cells per each height; *p* value, unpaired t test.

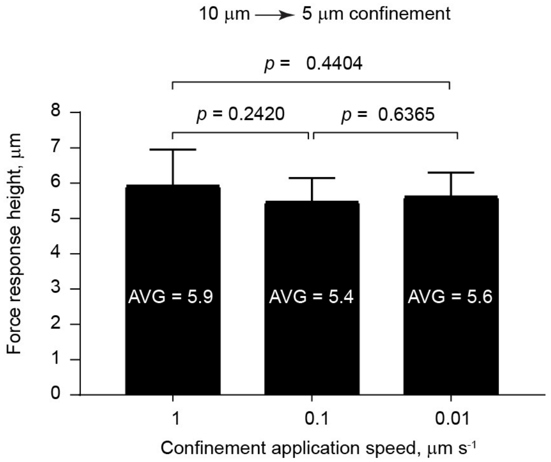


**Fig. S3.**

**The height for contractile force response is insensitive to cell deformation rate.**

Statistical analysis of the dependence between the confinement height at which cells display a robust contractile force response ($\Delta F > 15$ nN) to 10-to-5 μm confinement and the speed with which the confinement is applied. Mean ± SD; n = 10 cells per each experimental condition; *p* value, unpaired t test.

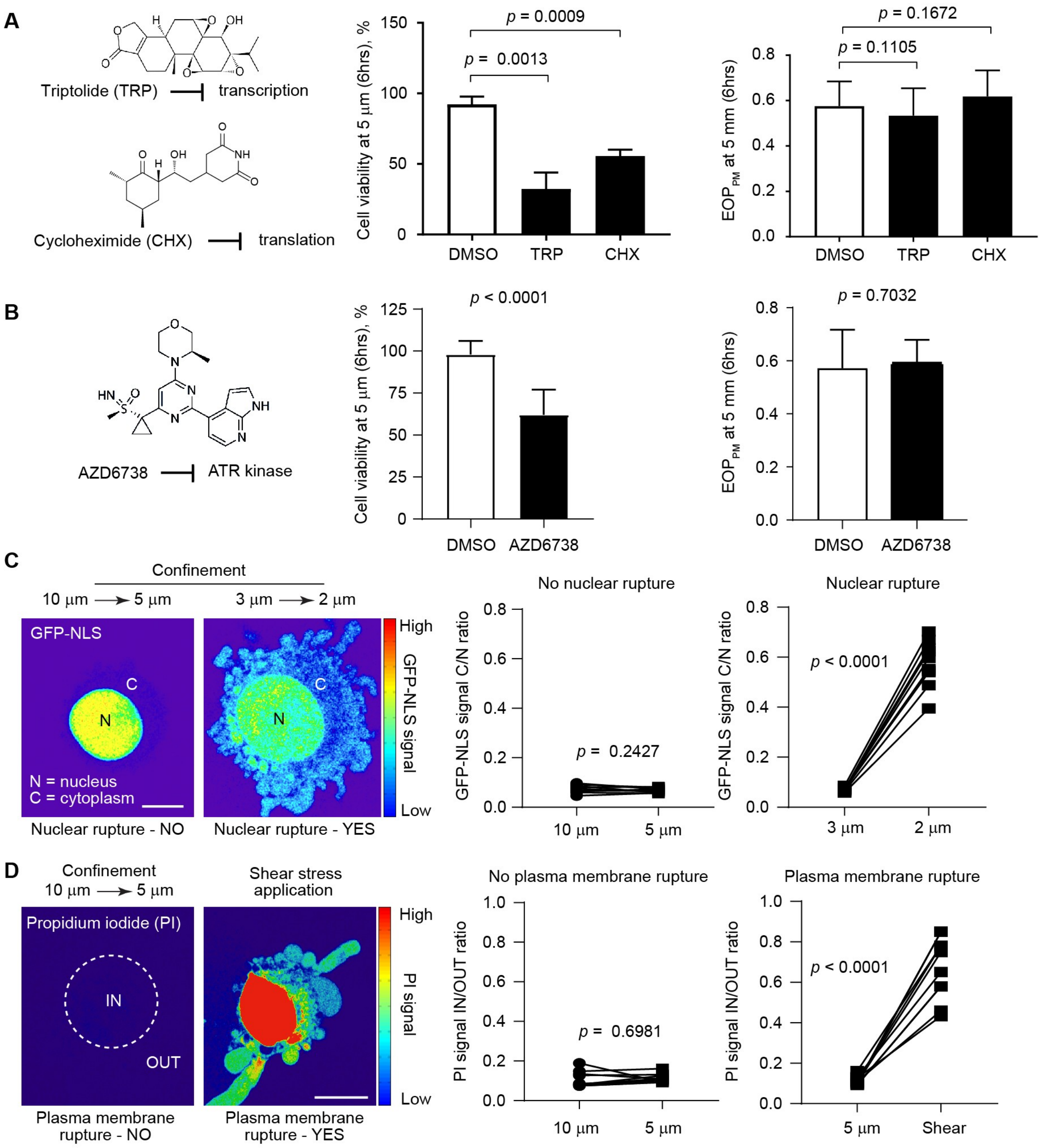


**Fig. S4.**

**Testing alternative mechanisms potentially implicated in the activation of myosin II-based contractility in response to precise confinement.**

**A:** Pretreatment of cells with pharmacological inhibitors of global transcription and translation does not interfere with the contractile cell output in the form of plasma membrane blebbing measured as $EOP_{PM}$ in cells under 5 μm confinement (right, mean ± SD; n = 35 cells per each experimental condition; *p* value, unpaired t test). The effect of the inhibitors on cell viability (middle) is used as a positive control for the inhibitors' activity (mean ± SD; n = 150 cells per each experimental condition; *p* value, unpaired t test).

**B:** Pretreatment of cells with a pharmacological inhibitor of ATR kinase does not interfere with the contractile cell output in the form of plasma membrane blebbing measured as $EOP_{PM}$ in cells under 5 µm confinement (right, mean ± SD; n = 40 cells per each experimental condition; *p* value, unpaired t test). The effect of the ATR kinase inhibitor on cell viability (middle) is used as a positive control for the inhibitor's activity (mean ± SD; n = 150 cells per each experimental condition; *p* value, unpaired t test).
**C:** Representative images of a GFP-labeled nuclear localization sequence (NLS) probe in cells subjected to 10-to-5 µm confinement (no nuclear rupture) *vs.* 3-to-2 µm confinement (nuclear rupture = positive control). Appearance of the GFP-NLS signal in the cytoplasm is a sign of nuclear membrane rupture. Measurements of the GFP-NLS signal subcellular redistribution in the same living cell subjected to 10-to-5 µm confinement (n = 10 different cells; *p* value, paired t test) or 3-to-2 µm confinement (n = 10 different cells; *p* value, paired t test) confirm that confining cells to 5 µm height does not rupture the nuclear membrane. Scale bar, 10 µm.
**D:** Representative images of Propidium iodide (PI) fluorescence in cells subjected to 10-to-5 µm confinement (no plasma membrane rupture) *vs.* shear stress (plasma membrane rupture = positive control). PI can enter the cell through nonspecific pores/wounds in the plasma membrane and becomes fluorescent after binding to cytoplasmic RNA or nuclear DNA molecules. Measurements of PI fluorescence levels in the same living cell subjected to 10-to-5 µm confinement (n = 10 different cells; *p* value, paired t test) or shear stress (n = 10 different cells; *p* value, paired t test), confirm that cell confinement *per se* does not rupture the plasma membrane. Scale bar, 15 µm.

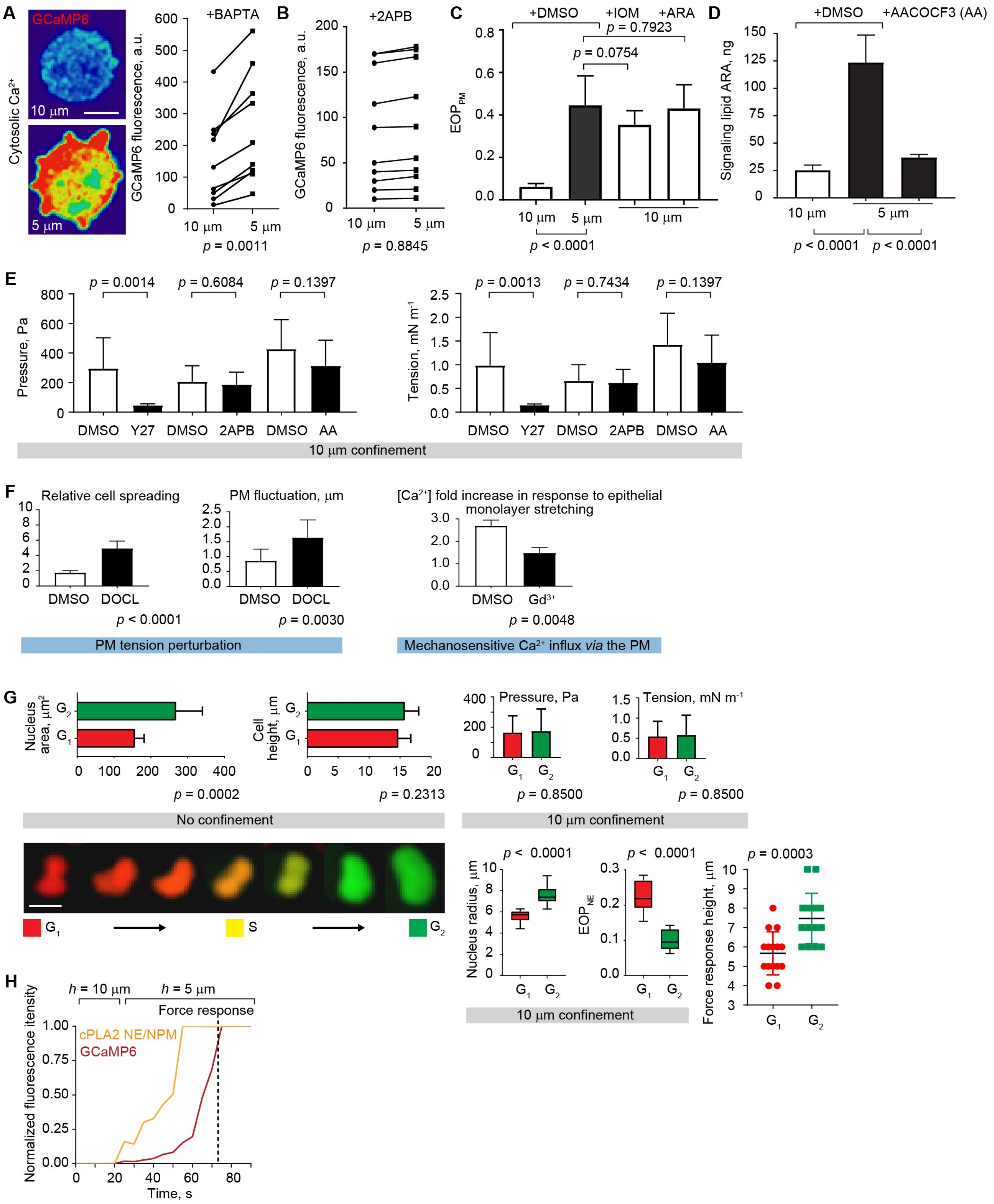


**Fig. S5.**

**Molecular pathways associated with nuclear and perinuclear ER membrane stretch become engaged at 5 µm confinement height without affecting cell behaviors at 10 µm height.**

**A:** Left, representative images illustrating that intracellular $Ca^{2+}$ concentration increases in response to 10-to-5 µm confinement even in the presence of BAPTA chelating extracellular $Ca^{2+}$. Right, measurements of cytoplasmic fluorescence intensity for the genetically encoded fluorescence $Ca^{2+}$ sensor GCaMP6 in the same living cell subjected to 10-to-5 µm confinement in the presence of BAPTA (n = 10 different cells; *p* value, paired t test). Scale bar, 10 µm.
**B:** Measurements of GCaMP6 fluorescence levels in the same BAPTA-treated living cell subjected to 10-to-5 µm confinement in the presence of 2APB inhibiting the stretch sensitive $Ca^{2+}$-channels InsP3R present on the perinuclear ER and nuclear membranes (n = 10 different cells; *p* value, paired t test).
**C:** Comparing the contractile cell output in the form of plasma membrane blebbing measured as $EOP_{PM}$ in cells under different degrees of spatial confinement in the presence of DMSO (control), ionomycin (IOM, artificially increasing the intracellular $Ca^{2+}$ level), or arachidonic acid (ARA, the signaling lipid-product of enzymatic activity of the nuclear membrane stretch-sensitive protein cPLA2). Mean ± SD; n = 35 cells per each experimental condition; *p* value, unpaired t test.
**D:** Biochemical measurements of ARA release in cell populations confined to 10 or 5 µm in the presence or absence of a cPLA2 inhibitor AACOCF3 (AA, n = 3 replicates; mean ± SD; *p* value, unpaired t test).
**E:** Measurements demonstrating that unlike perturbations that impact basal levels of actomyosin contractility (e.g. a Rho kinase/ROCK inhibitor Y27632/Y27), 2APB and AACOCF3 (AA) affecting the stretch sensitive $Ca^{2+}$-channels InsP3R on nuclear and perinuclear ER membranes and the nuclear membrane stretch-sensitive protein cPLA2 respectively do not change intracellular pressure and cortical tension estimated in cells at 10 µm confinement. Mean ± SD; n = 10 cells per each experimental condition; *p* value, unpaired t test.
**F:** Left, measurements demonstrating that deoxycholate (DOCL) does perturb plasma membrane (PM) tension as evidenced from DOCL's effect on relative cell spreading and fast fluctuations of the plasma membrane (n = 10 cells per each measurement; mean ± SD; *p* value, unpaired t test). Right, measurements of $Ca^{2+}$ entry into the cell upon stretching MDCK cell monolayers grown on flexible PDMS membranes in the presence or absence of $GdCl_3$ inhibiting stretch-sensitive ion channels on the plasma membrane (n = 3 replicates; mean ± SD; *p* value, unpaired t test).
**G:** Upper left, quantifications of nuclear size and cell height illustrating that rounded nonadherent G1 (n = 10) and G2 (n =10) cells have a similar height, but different nuclear size at resting state (no confinement). Upper right, quantifications of intracellular pressure and cortical tension in G1 (n = 10) *vs.* G2 (n = 10) cells illustrating that the cell cycle stage *per se* does not affect basal levels of actomyosin contractility that controls cortical cell tension and intracellular pressure at 10 µm confinement. Lower left, naturally occurring dynamic changes in nuclear shape and size accompanying cell cycle progression in the same live HeLa-Kyoto cell expressing the FUCCI fluorescent cell cycle reporter system and growing in a Petri dish. Lower right, quantifications revealing differences in nuclear size (radius) and folding ($EOP_{NE}$) in G1 (n = 10) *vs.* G2 (n = 10) cells under 10 µm confinement, and differential height-dependence of contractile force response in G1 (n = 15) *vs.* G2 (n = 15) cells. Mean ± SD; *p* value, unpaired t test. Scale bar, 10 µm.
**H**: A representative graph illustrating that an increase in cytosolic $Ca^{2+}$ and the accumulation at the nuclear envelope of the nuclear envelope stretch-sensitive enzyme cPLA2 temporally precede contractile force response measured with the flat AFM. This phenomenon is observed in almost 100% of analyzed cases (n = 30 cells).

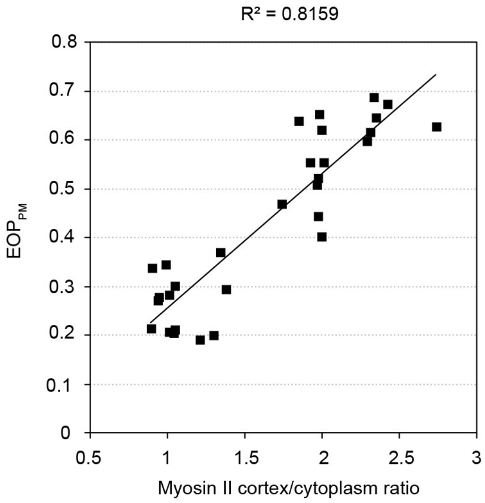


**Fig. S6.**

**Interdependence between the degree of plasma membrane blebbing and myosin II accumulation at the cortex in human fibrosarcoma cells maneuvering through 3D cell derived matrices.**

Correlation between the cell blebbing index measured as the excess of plasma membrane perimeter and cortical accumulation of myosin II in HT1080 cells imaged within the 3D dermal fibroblast-derived cell matrix (n = 30 cells).

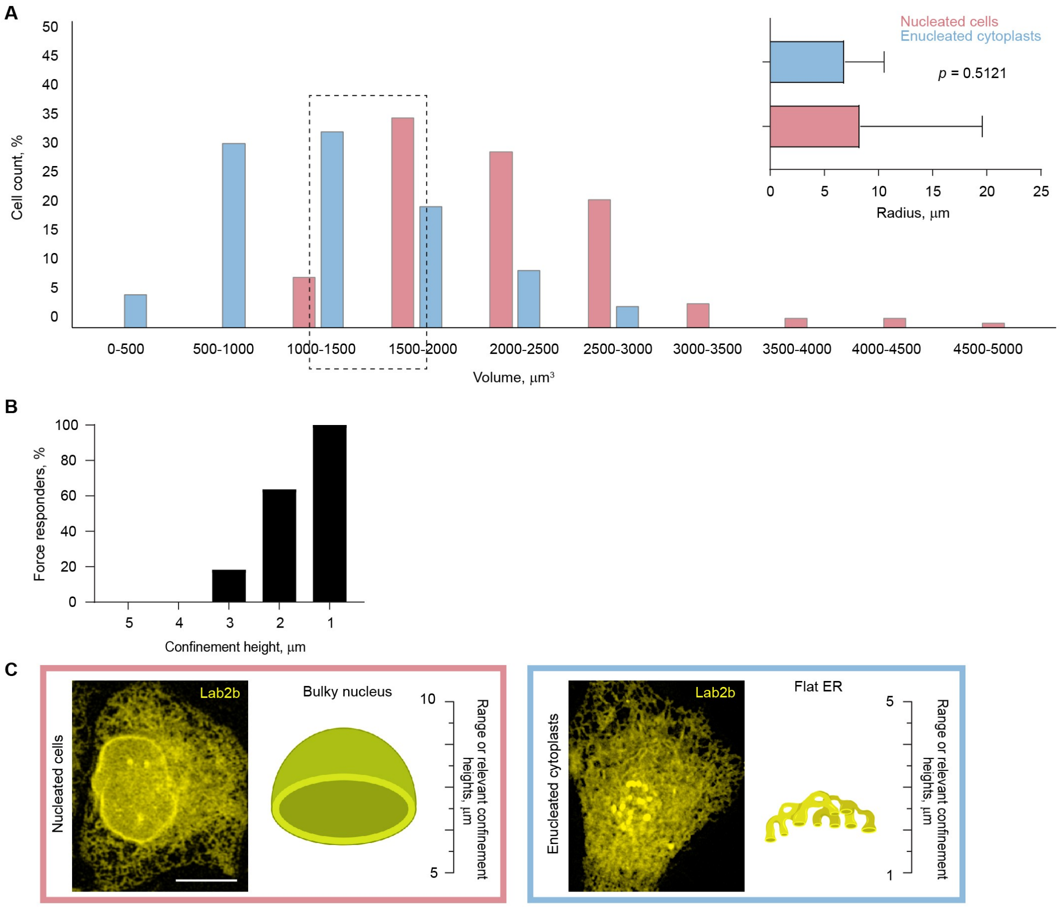


**Fig. S7.**

**Rounded enucleated cytoplasts possess a wide range of sizes, and those matching nucleated cell sizes are still incapable of measuring confinement height within the range of 10 to 5 µm.**

**A:** Large-scale statistical analysis of enucleated cytoplast size (n = 150) *vs*. nucleated cell size (n = 100, *p* value, unpaired t test).

**B:** Unlike rounded nucleated cells upon centrifugation, rounded enucleated cytoplasts with a similar size (~ 15 µm in diameter) do not respond to 10-to-5 µm confinement and begin measuring confinement height at significantly lower heights with a strong preference to 1 µm confinement (n = 25 cytoplasts with diameter ≥ 15 µm).

**C:** Representative images of Lap2B-GFP subcellular distribution illustrating that both nucleated cells and enucleated cytoplasts preserve an elaborate system of ER membranes that could take over the ruler function of the nucleus at significantly lower confinement heights. Scale bar, 10 µm.

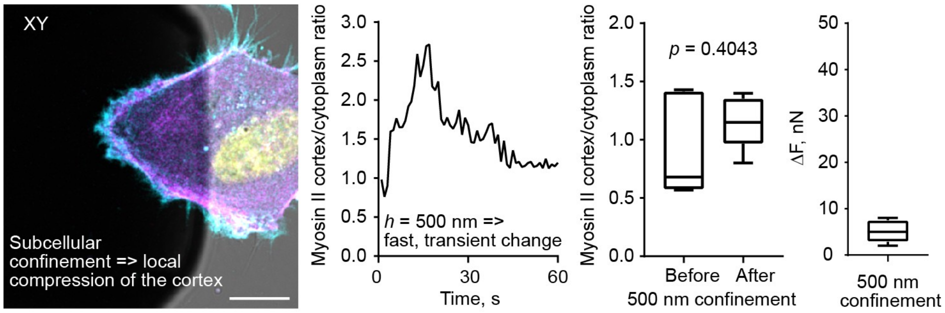


**Fig. S8.**

**The cortex compression in the lamellar region of the cell devoid of bulky cytoplasmic organelles.**

Left, an image illustrating the position of the flat cantilever to achieve a local compression of the cortex in the lamellar region of the cell (cyan, F-actin; magenta, myosin II; yellow, nucleus; scale bar, 10 µm), and quantification of the fast and transient increase in cortical myosin II signal upon strong (500 nm height confinement) compression of the cell cortex (the rapid and transient nature of the myosin II signal increase followed by decline to the initial level in response to local 500 nm compression of the lamellar region was observed in almost 100 % of analyzed cells (n = 15)). Right, statistical analyses of cortical accumulation of myosin II before and 5 minutes after 500 nm confinement of the cortex and force readouts after 500 nm confinement of the cortex (n = 10 cells; mean ± SD; *p* value, unpaired t test).

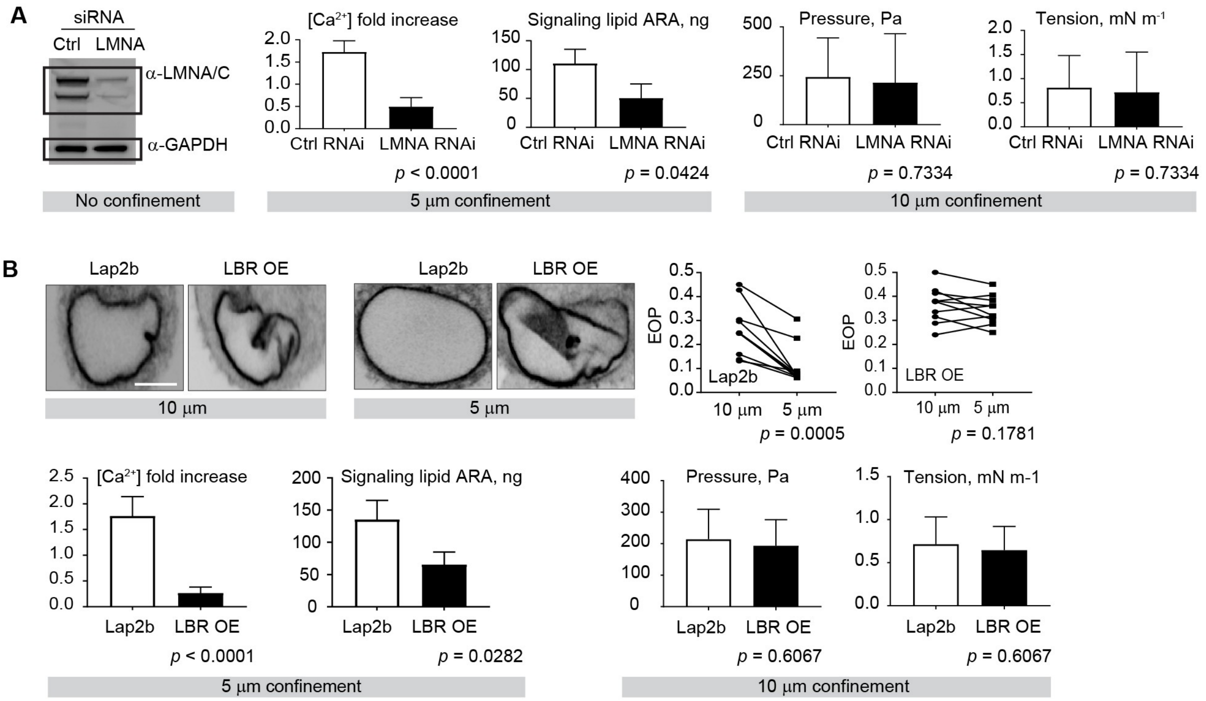


**Fig. S9.**

**Experimental perturbations targeting the ruler function of the nucleus do not affect basal contractility levels at 10 µm height, but significantly impact the signaling upstream of actomyosin activity at 5 µm confinement height.**

**A:** Left, nuclear LMNA knockdown efficiency in HeLa-Kyoto cells revealed with Western blotting. Middle, quantifications illustrating the effect of nuclear LMNA knockdown on the increase of intracellular $Ca^{2+}$ concentration and production of the signaling lipid ARA in response to 5 µm confinement (n = 3 replicates per each experimental condition; mean ± SD; *p* value, unpaired t test). Right, quantifications of intracellular pressure and cortical tension in control (n = 10) *vs.* LMNA depleted (n = 10) cells illustrating that LMNA depletion does not affect basal levels of actomyosin contractility that controls cortical cell tension and intracellular pressure at 10 µm confinement. Mean ± SD; *p* value, unpaired t test.

**B:** Top row, representative images of nuclei in control Lap2b-GFP expressing cells and LBR-GFP overexpressing (OE) cells at 10 vs. 5 µm confinement and quantifications of the degree of nuclear envelope folding measured as the excess of perimeter for the nuclear envelope ($EOP_{NE}$) in the same single live cell upon 10-to-5 µm confinement for Lap2b-GFP expressing cells and LBR-GFP OE cells (n = 10 different cells per each experimental condition; *p* value, paired t test). Bottom row left, quantifications illustrating the effect of LBR-GFP OE on the increase of intracellular $Ca^{2+}$ concentration and production of the signaling lipid ARA in response to 5 µm confinement (n = 3 replicates per each experimental condition; mean ± SD; *p* value, unpaired t test). Bottom row right, quantifications of intracellular pressure and cortical tension in control Lap2b-GFP expressing cells (n = 10) *vs.* LBR-GFP OE cells (n = 10) illustrating that LBR-GFP OE does not affect basal levels of actomyosin contractility that controls cortical cell tension and intracellular pressure at 10 µm confinement. Mean ± SD; *p* value, unpaired t test. Scale bar, 10 µm.

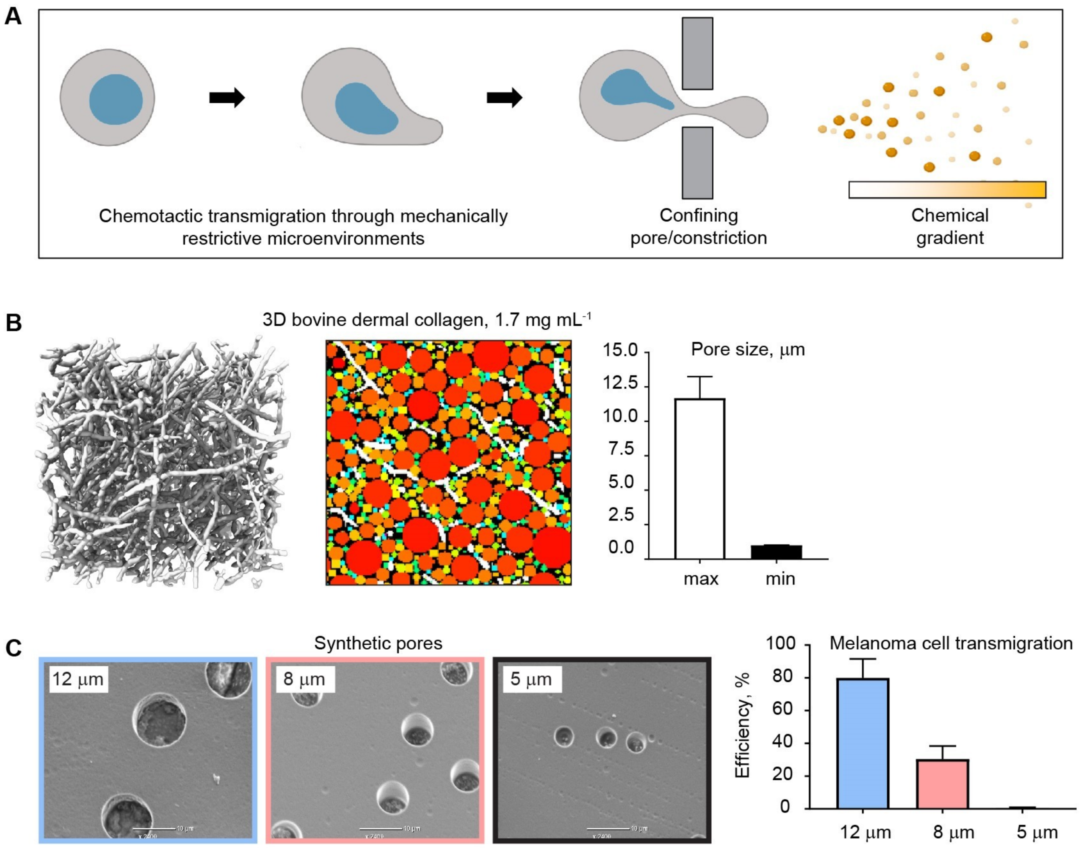


**Fig. S10.**

**Chemotactic transmigration of cells through mechanically restrictive pores and constrictions in 3D collagen gels and synthetic polycarbonate membranes.**

**A:** A simplified cartoon illustrating cell and nuclear squeezing through a confining opening in the extracellular environment in response to a gradient of chemotactic molecules, the situation that can be modelled experimentally using the Boyden chamber transwell assay.

**B:** A representative 3D super-resolution image of the dermal collagen lattice and an example of computer vision-aided pore size analysis by filling 3D spheres in between the collagen fibers in the skeletonized 3D super-resolution image of the collagen lattice. The measurements show that the 3D collagen lattices represent an environment with pore sizes ranging from ~ 1 µm (minimum size) to 10 µm (maximum size) in diameter.

**C:** Scanning electron microscopy images of polycarbonate membranes with 12, 8, and 5 µm pores in diameter, and quantification of the percentage of human melanoma cells A375 able to chemotactically transmigrate through the differentially-sized pores. The analysis shows that 5 µm pores are impenetrable for melanoma cells, 12 µm pores do not restrict cell migration, while 8 µm pores are rate-limiting to cell migration and thus ideally suited for the study of cell transmigration through mechanically restrictive environments.

Table S1. **Description of chemicals used in the mini-screen**

| **Pharmacological perturbations affecting plasma membrane (PM) tension & extracellular $Ca^{2+}$ influx** | | | **Pharmacological perturbations affecting stretch-sensitive proteins of the nuclear envelope (NE)/perinuclear ER & intracellular $Ca^{2+}$ homeostasis** | | |
|---|---|---|---|---|---|
| Drug name | Drug target/effect | Reference papers | Drug name | Drug target/effect | Reference papers |
| **Deoxy cholate (DOCL)** | Reduces PM tension | Raucher D, Sheetz MP. Cell spreading and lamellipodial extension rate is regulated by membrane tension. *J Cell Biol.* 2000 Jan 10;148(1):127-36.<br><br>Raucher D, Sheetz MP. Membrane expansion increases endocytosis rate during mitosis. *J Cell Biol.* 1999 Feb 8;144(3):497-506. | **AACOCF3 (AA)/ PACOCF3 (PA)** | Inhibits $Ca^{2+}$-dependent, NE tension-sensitive enzyme $cPLA_2$ producing the signaling lipid arachidonic acid (ARA) known to activate actomyosin contractility | Ackermann EJ, Conde-Frieboes K, Dennis EA. Inhibition of macrophage Ca(2+)-independent phospholipase A2 by bromoenol lactone and trifluoromethyl ketones. *J Biol Chem.* 1995 Jan 6;270(1):445-50.<br><br>Enyedi B, Jelcic M, Niethammer P. The Cell Nucleus Serves as a Mechanotransducer of Tissue Damage-Induced Inflammation. *Cell.* 2016 May 19;165(5):1160-1170.<br><br>Peters-Golden M, Song K, Marshall T, Brock T. Translocation of cytosolic phospholipase A2 to the nuclear envelope elicits topographically localized phospholipid hydrolysis. *Biochem J.* 1996 Sep 15;318 ( Pt 3):797-803.<br><br>Schievella AR, Regier MK, Smith WL, Lin LL. Calcium-mediated translocation of cytosolic phospholipase A2 to the nuclear envelope and endoplasmic reticulum. *J Biol Chem.* 1995 Dec 22;270(51):30749-54.<br><br>Gong MC, Fuglsang A, Alessi D, Kobayashi S, Cohen P, Somlyo AV, Somlyo AP. Arachidonic acid inhibits myosin light chain phosphatase and sensitizes smooth muscle to calcium. *J Biol Chem.* 1992 Oct 25;267(30):21492-8.<br><br>Feng J, Ito M, Kureishi Y, Ichikawa K, Amano M, Isaka N, Okawa K, Iwamatsu A, Kaibuchi K, Hartshorne DJ, Nakano T. Rho-associated kinase of chicken gizzard smooth muscle. *J Biol Chem.* 1999 Feb 5;274(6):3744-52.<br><br>Katayama T, Watanabe M, Tanaka H, Hino M, Miyakawa T, Ohki T, Ye LH, Xie C, Yoshiyama S, Nakamura A, Ishikawa R, Tanokura M, Oiwa K, Kohama K. Stimulatory effects of arachidonic acid on myosin ATPase activity and contraction of smooth muscle via myosin motor domain. *Am J Physiol Heart Circ Physiol.* 2010 Feb;298(2):H505-14. |
| **Gadolinium (III) chloride ($Gd^{3+}$)/ The peptide GsMTx4 from the tarantula venom** | Inhibits mechanosensitive ion channels enabling $Ca^{2+}$ influx *via* the PM | Gudipaty SA, Lindblom J, Loftus PD, Redd MJ, Edes K, Davey CF, Krishnegowda V, Rosenblatt J. Mechanical stretch triggers rapid epithelial cell division through Piezo1. *Nature.* 2017 Mar 2;543(7643):118-121.<br><br>Miyamoto T, Mochizuki T, Nakagomi H, Kira S, Watanabe M, Takayama Y, Suzuki Y, Koizumi S, Takeda M, Tominaga M. Functional role for Piezo1 in stretch-evoked $Ca^{2+}$ influx and ATP release in urothelial cell | **2APB/ Xestospongin C (Xesto)** | Blocks stretch-activated inositol triphosphate receptors (InsP3Rs) liberating $Ca^{2+}$ from membranes of the NE/perinuclear ER and provoking actomyosin | Kim TJ, Joo C, Seong J, Vafabakhsh R, Botvinick EL, Berns MW, Palmer AE, Wang N, Ha T, Jakobsson E, Sun J, Wang Y. Distinct mechanisms regulating mechanical force-induced $Ca^{2+}$ signals at the plasma membrane and the ER in human MSCs. *Elife.* 2015 Feb 10;4:e04876.<br><br>Solanes P, Heuzé ML, Maurin M, Bretou M, Lautenschlaeger F, Maiuri P, Terriac E, Thoulouze MI, Launay P, Piel M, Vargas P, Lennon-Duménil AM. Space exploration by dendritic cells requires maintenance of |

| | | | | | |
|---|---|---|---|---|---|
| | | cultures. *J Biol Chem.* 2014 Jun 6;289(23):16565-75. | | activity in a $Ca^{2+}$-dependent manner (e.g. *via* $Ca^{2+}$/calmodulin-dependent myosin light chain kinase/MLCK inhibited by the **ML-7** compound) | myosin II activity by IP3 receptor 1. *EMBO J*. 2015 Mar 12;34(6):798-810.<br><br>Cárdenas C, Liberona JL, Molgó J, Colasante C, Mignery GA, Jaimovich E. Nuclear inositol 1,4,5-trisphosphate receptors regulate local Ca2+ transients and modulate cAMP response element binding protein phosphorylation. *J Cell Sci.* 2005 Jul 15;118(Pt 14):3131-40.<br><br>Bootman MD, Fearnley C, Smyrnias I, MacDonald F, Roderick HL. An update on nuclear calcium signalling. *J Cell Sci.* 2009 Jul 15;122(Pt 14):2337-50. |
| **BAPTA** | Chelates $Ca^{2+}$ specifically outside the cell | Solanes P, Heuzé ML, Maurin M, Bretou M, Lautenschlaeger F, Maiuri P, Terriac E, Thoulouze MI, Launay P, Piel M, Vargas P, Lennon-Duménil AM. Space exploration by dendritic cells requires maintenance of myosin II activity by IP3 receptor 1. *EMBO J.* 2015 Mar 12;34(6):798-810. | **BAPTA-AM** | Chelates $Ca^{2+}$ specifically inside the cell | Kono T, Jones KT, Bos-Mikich A, Whittingham DG, Carroll J. A cell cycle-associated change in Ca2+ releasing activity leads to the generation of Ca2+ transients in mouse embryos during the first mitotic division. *J Cell Biol*. 1996 Mar;132(5):915-23. |

**Movie S1.**

Rounded nonadherent HeLa-Kyoto cells sense the difference between 10 and 5 μm confinement – spatially confining cells to 5 but not 10 μm height stimulates rapid recruitment of myosin II from the cytosol to the cortex. Left, a representative cell expressing Myh9-GFP (pseudocolored using the thermal LUT mode with hot and cold tones indicating high and low levels of Myh9-GFP fluorescence signal respectively) immediately after the 10 μm height is reached. Right, the same cell after 5 μm confinement. Time interval between consecutive images, 5 s. Scale bar, 5 μm.

**Movie S2.**

Rounded nonadherent HeLa-Kyoto cells undergo a discrete morphodynamic switch from a noncontractile cell cortex and filopodia formation to a highly contractile cortex and vigorous plasma membrane blebbing in response to 10-to-5 μm confinement. Left, a representative cell expressing Myh9-GFP (pseudocolored in magenta) and LifeAct-mCherry (pseudocolored in cyan) during 20-to-10 μm confinement. Right, the same cell during 10-to-5 μm confinement. A differential-interference-contrast (DIC) image of the confining flat microcantilever is shown in the grey channel. Time interval between consecutive images, 5 s. Scale bar, 5 μm.

**Movie S3.**

Rounded nonadherent HeLa-Kyoto cells expand their nuclei and unfold the nuclear envelope prior to the onset of the contractile cell response manifested in active plasma membrane blebbing during 10-to-5 μm confinement. The representative cell expresses Lap2b-GFP as a marker of the nuclear envelope (green) and is stained with the SiR-Actin probe to reveal F-actin (pseudocolored in magenta). The cell was confined from 20-to-10-to-5 μm. The video is artificially paused several times during the 10-to-5 μm transition to accentuate nuclear expansion-unfolding taking place before the cell retracts the very first rounded blebs. These structures represent passive bleb-like protrusions appearing upon physical squeezing of the cell (i.e. when intracellular fluid is pushed against the plasma membrane through pre-existing cracks in the cell cortex). Our additional observations (data not shown) reveal that such "passive" blebs almost instantaneously accumulate submembranous cortical F-actin. The cortex of the "passive" blebs gets retracted by active myosin II motors initiating a continuous active blebbing. "Active" blebs are more numerous, often of a larger length, and significantly smaller in width compared to their "passive" counterparts. Time interval between consecutive images, 5 s. Scale bar, 5 μm.

**Movie S4.**

Rounded nonadherent, enucleated HeLa-Kyoto cells (cytoplasts) do not sense the difference between confinement heights relevant to the contractile response in nucleated cells. Left, a representative nucleated cell expressing Myh9-GFP (pseudocolored in magenta) and LifeAct-mCherry (pseudocolored in cyan) that survived the treatments and centrifugation protocol required for enucleated cytoplast generation. Right, a representative enucleated cytoplast. The video starts from depicting a nonconfined state of these cells followed by 20-to-10-to-5 μm confinement. Note, despite its smaller size (height), the nucleated cell that survived the cytoplast generation protocol initiates the sustained, active contractile response (cortical accumulation of myosin II and vigorous plasma membrane blebbing) right before the confining cantilever stops at 5 µm during the movement from 10 to 5 µm. However, the enucleated cytoplast displays only a passive response in the form of transient bleb-like protrusions generated due to physical squeezing of the cytoplast during the 10-to-5 µm confinement. Time interval between consecutive images, 5 s. Scale bar, 5 µm.

**Movie S5.**

Compression of the cell cortex down to 500 nm confinement height yields only local and highly transient bleb-like protrusions without any global effect on the levels of actomyosin contractility. Left, a representative spread adherent HeLa-Kyoto cell expressing Myh9-GFP (pseudocolored in magenta), LifeAct-mCherry (pseudocolored in cyan) and additionally stained with DAPI (pseudocolored in yellow). A differential-interference-contrast (DIC) image of the confining flat microcantilever is shown in the grey channel. The precise position of the cantilever was adjusted such that only the lamellar but not organelle-rich compartment is engaged in the local cell compression. Right, a time-lapse video of the cell responding to strong local cortical compression (500 nm confinement). Time interval between consecutive images, 5 s. Scale bar, 10 µm.

**Movie S6.**

Induction of the global sustained contractile cell response to local nuclear but not cortex/cytoplasm confinement. The video depicts representative spread adherent HeLa-Kyoto cells expressing Myh9-GFP (pseudocolored in magenta), LifeAct-mCherry (pseudocolored in cyan) and additionally stained with DAPI (pseudocolored in yellow) to reveal the nuclear compartment. A differential-interference-contrast (DIC) image of the confining flat microcantilever is shown in the grey channel. The precise position of the cantilever was adjusted such that only the nuclear compartment of the lower cell and the lamellar region of the upper cell are engaged in the local confinement. Confinement height, 2 µm. Note, while local cortex/cytoplasm confinement produces local, short-lived bleb-like protrusions (passive response to physical compression), local nuclear-specific confinement results in a global, sustained increase in cell contractility even in the non-confined cell region that switches to this state with a delay (activation signal propagation). Time interval between consecutive images, 5 s. Scale bar, 10 µm.
